# Stress drives plasticity in leaf maturation transcriptional dynamics

**DOI:** 10.1101/2025.02.24.639183

**Authors:** Joseph Swift, Xuelin Wu, Jiaying Xu, Tanvi Jain, Natanella Illouz-Eliaz, Joseph R. Nery, Joanne Chory, Joseph R. Ecker

## Abstract

Leaf development is dynamic, adapting to environmental stress to optimize resource use. For example, in response to drought, *Arabidopsis* restricts leaf growth to enhance water use efficiency. While this plastic response is well described at the physiological level, the underlying transcriptional changes that facilitate adjustments in leaf development in response to stress remain poorly described. By constructing a ∼1 million single-nuclei transcriptome atlas, we demonstrate that drought stress limits leaf growth by advancing transcriptional responses related to maturation and aging. Notably, we find that these transcriptional changes scale with environmental stress, and help explain how shoot size can decline proportionally with stress intensity. Leveraging these insights, we increased leaf growth under stress by cell-type targeted upregulation of *FERRIC REDUCTION OXIDASE 6* in the mesophyll.

## Main Text

Plant leaf growth progresses through ordered stages of development, yet unlike mammalian development, it exhibits a remarkable degree of plasticity by responding dynamically to environmental cues. For instance, upon encountering drought stress, the model plant *Arabidopsis thaliana* will limit the size of growing leaves and induce senescence in older ones (*1-3*). Referred to as the ‘stress avoidance’ strategy (*4*), this phenotypic change leads to a compact plant stature that is more water use efficient, and helps plants survive when drought conditions arise (*4, 5*). The gene regulatory responses that drive such plasticity are not well understood. Since leaf growth is underpinned by the sequential expression of genes that direct cellular proliferation, expansion and senescence (*6*), changing the timing or induction levels of such genes may be one way in which drought signaling can impact the course of leaf development. Indeed, drought stress has been shown to downregulate genes associated with leaf growth rate (*7*), as well as to induce cell cycle exit (*8*). However, a global picture of how the transcriptional events that drive leaf development are reshaped by drought stress remains incomplete.

To address this, we characterized the genome-wide leaf maturation transcriptional changes among *Arabidopsis* cell types and explored how their expression dynamics changed as drought intensified. Across cell types, we found that drought stress largely promotes leaf maturation signals, and we found that this coincides with a change in hormone signaling associated with leaf development. Furthermore, we determined that maturation associated genes adjust their expression levels in a dose-dependent manner, and posit that such responses explain how *Arabidopsis* can limit shoot size in proportion to stress intensity. Our resulting transcriptional atlas led us to identify *FERRIC REDUCTION OXIDASE 6* (FRO6) within the mesophyll cell type, and by perturbing its cell-type specific expression pattern, we increased leaf growth responses under drought stress.

## Results

### Dynamic transcriptional cell states underlie leaf maturation

An *Arabidopsis* rosette contains leaves at various stages of development. To investigate the transcriptional changes associated with leaf maturation, we took advantage of this variation by harvesting leaves across 15 distinct leaf developmental stages displayed by mature rosettes. Moreover, we sampled each of these leaf stages over a 9 day time course. By these means, we collected a total of 647 leaves at various developmental and temporal ages (**Fig. 1a**). To profile cell-type specific transcriptomes of each individual leaf, we employed sci-RNA-seq3 (*9*). This technique enabled us to uniquely barcode and sequence each leaf to single-nuclei resolution (**Fig. 1a**). Using this data, we constructed an *Arabidopsis* leaf single-nuclei atlas comprising 264,183 nuclei (**Fig. 1b**, **Table S1**). By relying on validated cell identity markers, we annotated the majority of clusters within our atlas and identified nine leaf cell types (**Fig. S1**, **Table S2**). This atlas can be explored through an interactive browser (**Fig. S2**).

**Fig. 1:**
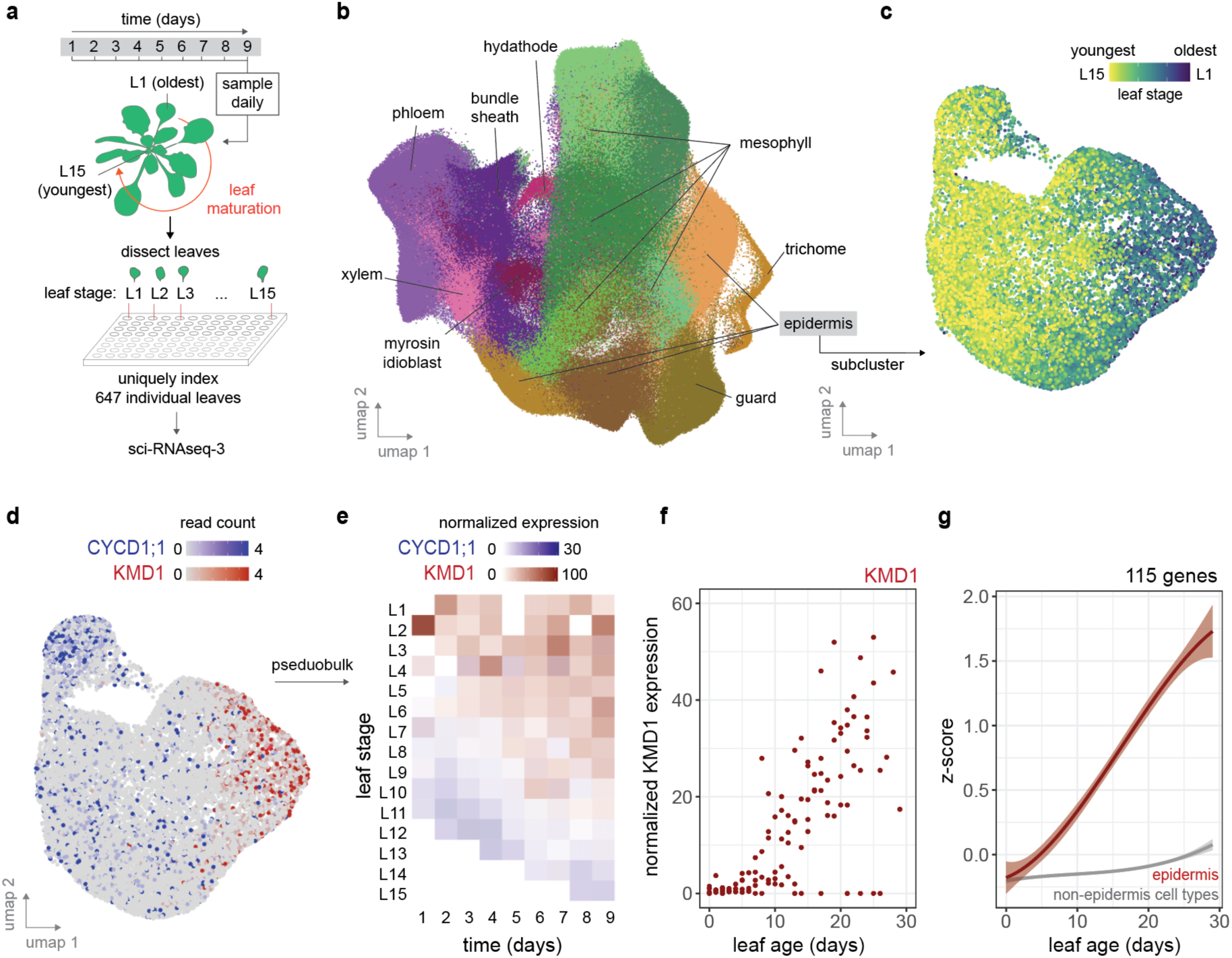
Transcriptional states of *Arabidopsis* cell types change during leaf maturation. **(a)** 647 individual leaves sourced across 15 leaf stages and 9 days of growth were sequenced to single nuclei resolution using sci- RNA-seq3. **(b)** A leaf transcriptional atlas was assembled from the resulting 264,183 nuclei, and 9 leaf cell types were identified using known marker genes. **(c)** Epidermal nuclei subclustered and colored by their respective leaf stage. **(d)** Expression patterns of *CYCD1;1* and *KMD1* within the epidermal sub-cluster. **(e)** Pseudo-bulked expression profiles of *CYCD1;1* and *KMD1* in the epidermis cell-type, arranged by leaf stage and day of sampling. **(f)** Expression profile of *KMD1* across leaf age (n = 118 averaged replicates). **(g)** z-score normalized expression trend of 115 genes upregulated specifically within the epidermis cell type during leaf maturation (curve fit using quadratic model, 99% confidence interval indicated).

We observed a dynamic shift in cell transcriptional states as leaves grew and matured. This was evident through the unsupervised sub-clustering of each cell type, which resulted in nuclei organizing by their leaf development stage (**Fig. 1c**, **Fig. S3**). For example, by sub-clustering the epidermal cell type, we found that the cell cycle gene *CYCLIN D1;1* (*CYCD1;1*) was expressed in the youngest leaves, indicating that nuclei were sourced from leaves undergoing cellular proliferation (*10*) (**Fig. 1d**). Conversely, we found the negative regulator of cytokinin signaling *KISS ME DEADLY 1* (*KMD1*) expressed among nuclei sourced from the oldest leaves (*11*). Pseudo- bulking epidermal nuclei by their respective leaf stage and sampling time point provided further insight into the transcriptional dynamics of leaf maturation. For instance, we discovered that *KMD1* expression increased as the leaf stage progressed (**Fig. 1e**), and that its expression significantly rose as each leaf stage matured over real time (adj. *p* = 1.92 × 10^-52^, linear model) (**Fig. 1e**). Similarly, when we calculated a leaf’s age based on its developmental stage and the time point assayed, we found that *KMD1*’s expression increased as leaves aged (**Fig. 1f, Fig. S4**). We noted that *CYCD1;1* expression was downregulated as *KMD1* expression was upregulated (**Fig. S4**).

We identified similar expression trends genome-wide and across cell types, with hundreds of genes that significantly increased or decreased in their expression as leaves matured (adj. *p* < 0.01, linear model), including established leaf development genes (**Fig. S3**). Many of these patterns were specific to certain cell types (**Table S3**). For instance, we found 115 genes that were specifically upregulated in the epidermis cell type as leaves aged, holding enriched Gene Ontology (GO) terms such as ‘response to biotic stress’ and ‘response to auxin stimulus’ (adj. *p* < 8.8 × 10^-3^) (**Fig. 1g**). Among these were the auxin signaling gene *INDOLE-3-ACETIC ACID INDUCIBLE 29* (*IAA29*) and defense related transcription factor *WRKY DNA-BINDING PROTEIN 38* (*WRKY38*), indicating that auxin and defense signaling coordination in leaf senescence occurs in the leaf epidermis (*12*).

### Drought stress advances leaf maturation transcriptional dynamics

When drought conditions arise *Arabidopsis* leaf growth is attenuated. To understand the transcriptional changes underlying this plastic response, we examined how cell-type specific leaf maturation transcriptional dynamics changed during the onset of drought. To achieve this, we subjected vermiculite grown *Arabidopsis* rosettes to drought stress by withholding water for nine days, which decreased the available water content (WC) within the vermiculite from 100 % to 21 %. As expected, this treatment resulted in smaller leaf sizes and reduced shoot biomass (**Fig. 2a, 2b, Fig. S5, Table S4**). We employed sci-RNA-seq3 to assay gene expression responses among individual leaves at single nuclei resolution, adding 173,731 nuclei from 579 leaves to our leaf atlas, which represents 15 leaf stages across nine time points of the drought stress assay (**Fig. S1**).

**Fig. 2:**
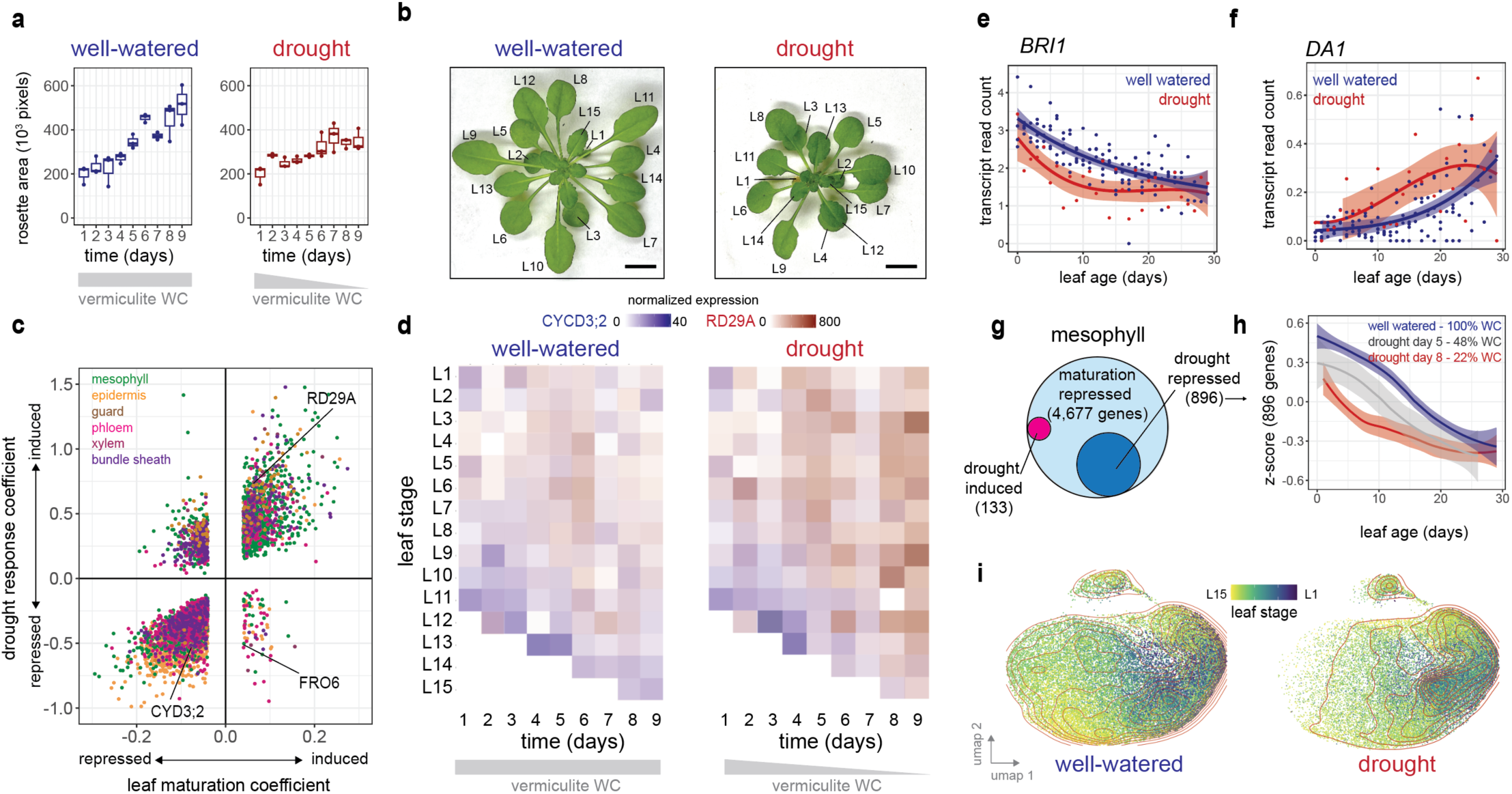
Drought stress promotes leaf maturation transcriptional dynamics. **(a)** Whole rosette area of 29- day old *Arabidopsis* plants grown under well-watered conditions or subjected to drought stress by withholding water over 9 days (n = 2-3, median indicated, *p* = 3.4 × 10^-3^, ANCOVA). **(b)** Images of well-watered or drought stressed rosettes on the 9^th^ day (leaf stage indicated; bar represents 1 cm). **(c)** Induction or repression of leaf maturation genes found to be differentially expressed in response to drought stress (linear model coefficient indicated, adj. *p* <0.01, color indicates cell-type). **(d)** Pseudo-bulked mesophyll transcriptional profiles of *CYCD3;2* and *RD29A* expression across leaf stages and over time under well-watered or drought conditions. **(e)** *BRI1* and **(f)** *DA1* expression across leaf age within the mesophyll cell type under well-watered conditions (100% WC) and drought conditions (22 - 28% WC, line fit using quadratic model, 95% confidence interval indicated). **(g)** The number of genes repressed during leaf maturation in the mesophyll cell type, and the subset that is responsive to drought stress. **(h)** Normalized z-score expression trends of 896 leaf maturation genes under three levels of vermiculite water content (WC, 99 % confidence interval indicated). **(i)** Mesophyll nuclei sourced from well-watered or drought conditions were subclustered and colored by their respective leaf stage (red contour lines represent levels of equal point density; both UMAPs present 30,270 nuclei).

Across various cell types, we found that drought stress enhanced the expression of genes activated during leaf maturation, while simultaneously repressing those involved in cellular proliferation and expansion (**Fig. 2c, Fig. S4, Fig. S6**). For example, under well-watered conditions, the expression of *RESPONSE TO DESICCATION 29A* (*RD29A*) was induced in the mesophyll cell-type as leaves matured. However, as drought stress intensified, *RD29A* was precociously upregulated in younger leaves, coinciding with the repression of *CYCLIN D3;2* (*CYCD3;2*) (**Fig. 2d**). Similarly, genes that positively or negatively regulate leaf size – such as *BRASSINOSTEROID INSENSITIVE 1* (*BRI1*) (*13*) and *DA* (*DA1*) (*14*) respectively – were prematurely down- or up-regulated within the mesophyll in response to drought (**Fig. 2e, f**). This trend extended to other genes involved in leaf development, cell cycle, and photosynthesis (**Fig. S6**), and was observed genome-wide across cell types (**Table S5**). For instance, of the 4,677 genes significantly downregulated in the mesophyll during leaf maturation (adj. *p* < 0.01, linear model), drought prematurely repressed 896 (19 %) of these genes. The degree of repression coincided with decreasing vermiculite water content (**Fig. 2g, h**). In contrast, it was relatively rare for drought stress to act antagonistically by inducing genes that were repressed during leaf maturation, or vice versa (**Fig. 2c, 2g**). Notably, we found that drought stress caused nuclei from younger leaves to cluster more closely with those from older leaves (**Fig. 2i, Fig. S5**), suggesting that drought stress prompts to transcriptional cell states in young leaves to resemble those of older leaves.

### Drought stress impacts hormone signaling responses to promote leaf maturation

We hypothesized that drought stress might enhance the expression of leaf maturation genes by influencing the hormone signals involved in leaf development. To investigate this, we treated whole rosettes with eight different phytohormones for two hours before sequencing leaf tissue to single nuclei resolution (**Fig. S1**). Using this approach, we identified hundreds of differentially expressed genes responsive to specific hormone treatments in either mesophyll, epidermal, or vasculature cell-type classes (**Fig. 3a, 3b, Fig. S7**). We then compared these responsive genes with those we identified as differentially expressed during leaf maturation or drought onset (**Table S6**).

**Fig. 3:**
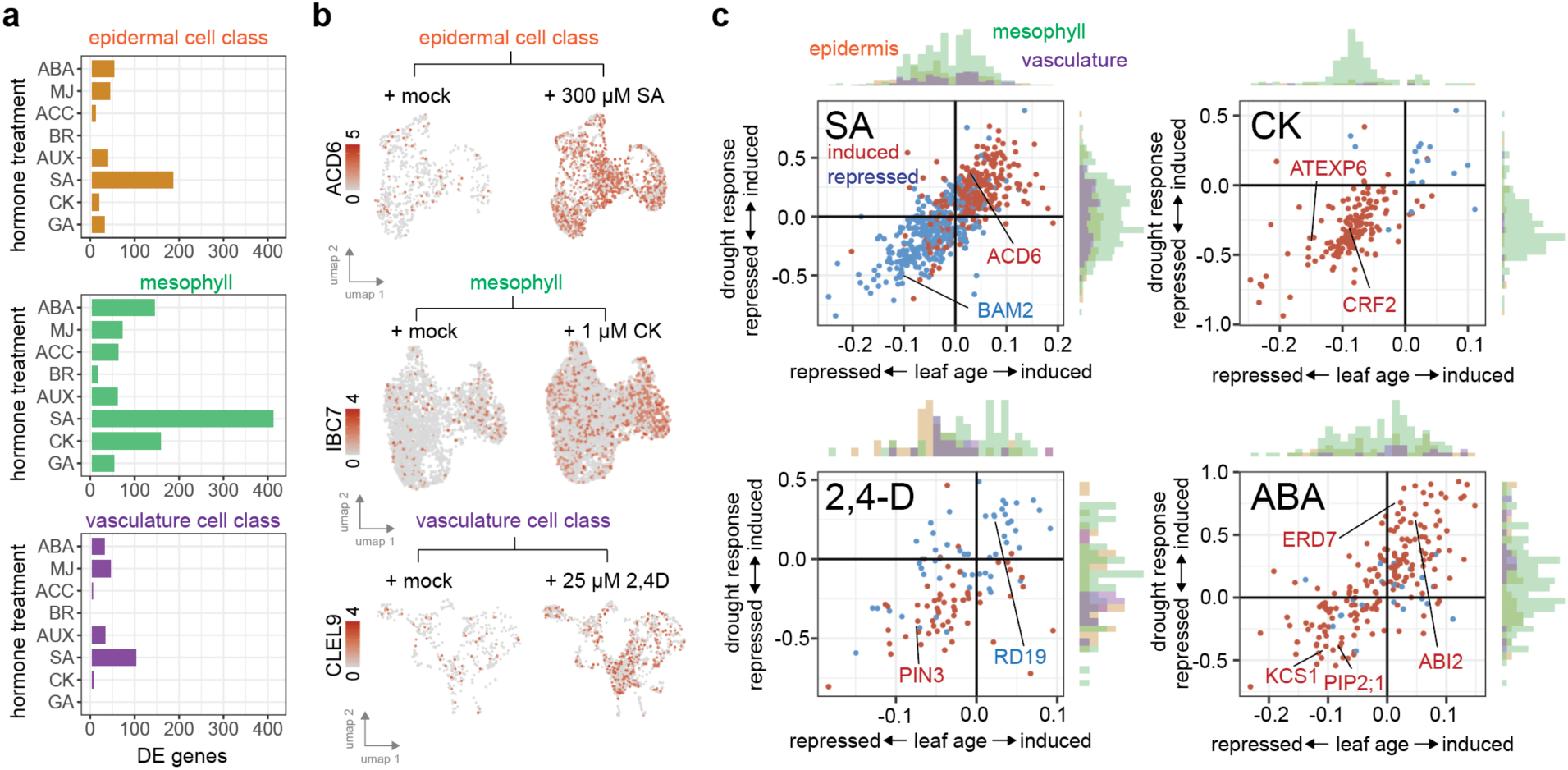
Hormone signals are linked to drought-induced changes to leaf maturation transcription dynamics. **(a)** Number of differentially expressed (DE) genes in either the mesophyll, the epidermal cell class (combining epidermis, guard and trichome cell types) or vasculature cell class (combining phloem, xylem, bundle sheath, hydathode, myrosin idioblast cell types) in response to transient treatment with either abscisic acid (ABA), methyl jasmonate (MJ), ethylene precursor 1-Aminocyclopropane-1-carboxylic acid (ACC), brassinollide (BR), synthetic auxin (2,4-D), salicylic acid (SA), cytokinin as *trans*-Zeatin (CK), and gibberellin (GA) (adjusted *p* < 0.1, likelihood-ratio test). **(b)** Single nuclei expression responses to exogenous hormone treatment of *ACD6*, *INDUCED BY CYTOKININ 7 (IBC7)* and *CLE-LIKE 9* (*CLEL9*) in epidermal, mesophyll and vasculature cell-type clusters respectively. **(c)** Induction (red) or repression (blue) of genes significantly responsive to one of four hormone treatments (adj. *p* < 0.1, likelihood ratio test), along with their respective induction or repression in response to leaf maturation or drought stress (adj. *p* < 0.01, linear model, axis units are coefficients of linear model). Histograms next to each axis show the cell-type class in which the hormone responsive gene was detected.

We found evidence that hormone signals promoting leaf maturation were further induced by drought stress in a cell class specific manner. For example, in accordance with salicylic acid’s (SA) role in advancing leaf maturation and senescence (*15*), we identified SA-induced genes significantly enriched among leaf maturation induced genes. We found these same genes were also drought induced (**Fig. 3c**). This included the leaf maturation regulator *ACCELERATED CELL DEATH 6* (*ACD6*) (*16*) within the epidermis cell class (**Fig. 3b, 3c**). Similarly, we found that SA-repressed genes enriched among leaf maturation repressed genes that were also drought repressed, including the leaf size regulator *BARELY ANY MERISTEM 2* (*BAM2*) (*17*) in the epidermal cell class (**Table S6**). Globally, genes within this overlap were enriched in stress and defense-related Gene Ontology (GO) terms (e.g. ‘defense response’ *p* = 4.0 × 10^-30^). Collectively, these data indicate that the onset of drought induces SA signaling to promote leaf maturation and senescence.

Additionally, we found evidence that hormone signals associated with leaf proliferation and expansion were repressed by drought stress in a cell type specific manner. For example, consistent with cytokinin’s (CK) role in promoting cellular proliferation (*10, 18*), we observed significant overlap between CK-induced genes and those downregulated both by leaf maturation and during drought onset (**Fig. 3c**). This enrichment was almost exclusively within the mesophyll cell type, with 60 % of members (82 genes) related to protein translation. These findings suggest that the onset of drought represses CK signaling to advance leaf maturation. Other hormone treatments (gibberellin (GA), brassinosteroid (BR), ethylene (ACC) and synthetic auxin (2,4-D)) elicited similar responses, but varied by the cell type classes they affected (**Fig. S8**).

We note that ABA-induced genes were enriched among leaf maturation induced genes that were also drought induced. In agreement with ABA’s role in signaling stress, these genes were over-represented in the ‘response to water deprivation’ GO term (*p* = 1.1 × 10^-22^). Curiously, we also found ABA-induced genes that were repressed during leaf maturation and drought stress. Among these genes were those whose upregulation might promote stress tolerance – such as 3-ketoacyl-CoA synthase 1 (*KCS1*) (*19*) in wax biosynthesis, and *PLASMA MEMBRANE INTRINSIC PROTEIN 2;1* (*PIP2;1*) (*20*) in water transport.

### Leaf maturation gene expression scales with stress intensity

The degree of drought severity may determine the degree to which leaf maturation expression patterns change. Supporting this hypothesis, we found evidence that transcriptional responses among rosette leaves changed proportionally as water availability declined (**Fig. 2h**). To investigate this effect further, we grew *Arabidopsis* seedlings for 11 days under eight increasing levels of ‘hard agar’ (HA) stress, a treatment that allows for more precise control of water potential compared to vermiculite drying in pots (*21*). As expected, as the level of HA stress intensified, we found shoot size declined in a dose-dependent manner with the intensity of stress (**Fig. 4a, 4b**). We profiled shoot cell-type transcriptomes under each HA dose at each time point tested using sci-RNA- seq3 (88 conditions). By these means, we sequenced a total of 456,008 nuclei and identified cell types by integrating nuclei with our rosette leaf atlas (**Fig. 4c**, **Fig. S1**).

**Fig. 4:**
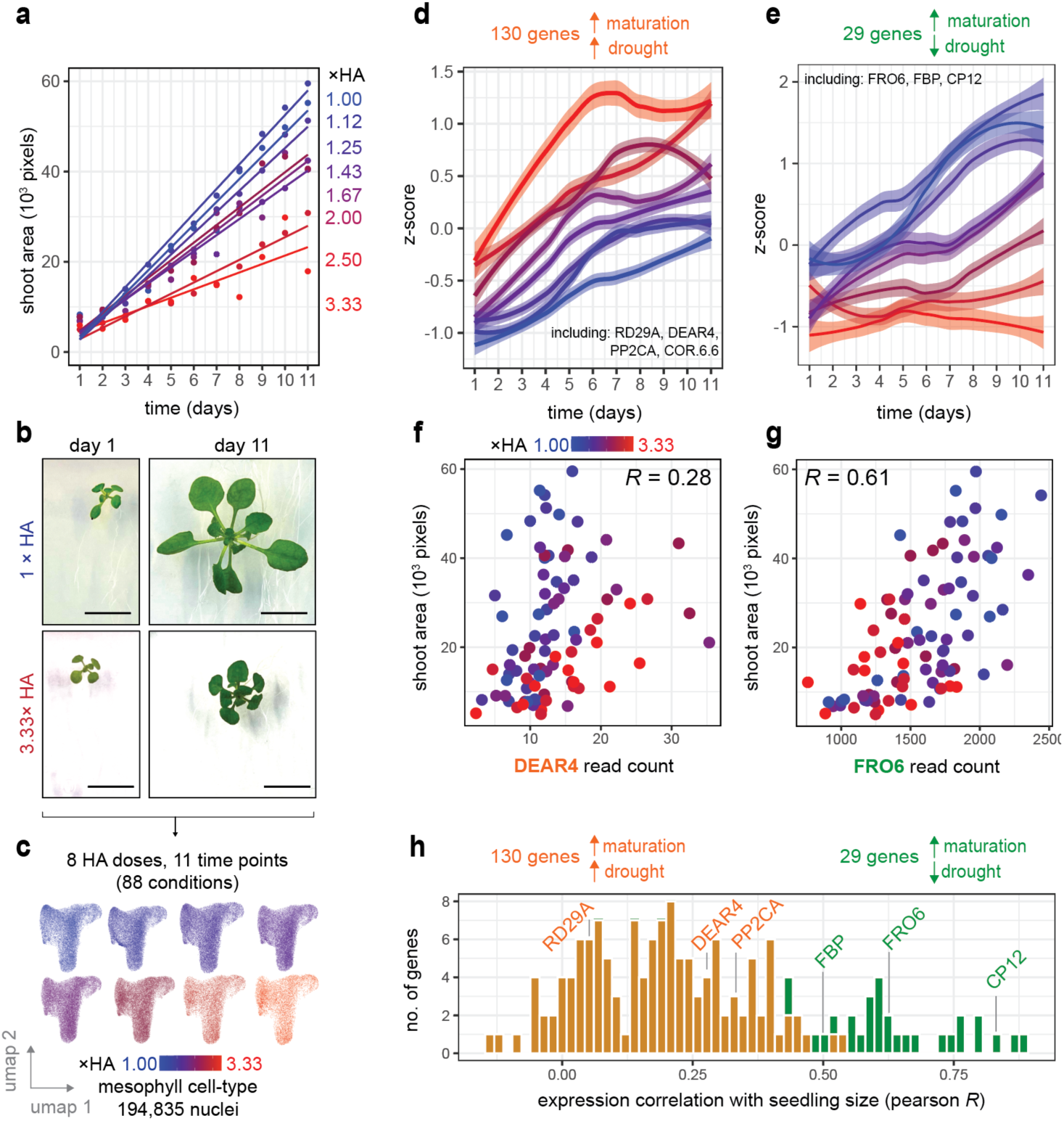
Dose-responsive transcriptional changes to hard agar (HA) stress are associated with shoot plasticity. **(a)** Shoot size of *Arabidopsis* seedlings grown for 11 days on eight different HA media doses. **(b)** Images of seedlings on the 1^st^ and 11^th^ day of 1× (no stress) or 3.3× HA stress treatment (bar represents 1 cm). **(c)** 595 individual shoots spanning the 11 days and 8 HA doses tested were sequenced to nuclei resolution using sci-RNA-seq3. Clusters present 194,835 nuclei from the mesophyll cell type, separated by the HA dose assayed. **(d)** Z-score plot of 130 genes whose expression in the mesophyll cell-type was induced during leaf and shoot maturation, and where HA stress induced expression levels further (line fit using locally estimated scatterplot smoothing, 99% confidence interval indicated). **(e)** Z-score plot of 29 genes whose expression in the mesophyll cell-type was induced during leaf and shoot maturation, and where HA stress acted ‘antagonistically’ by repressing gene expression (line fit using locally estimated scatterplot smoothing, 99% confidence interval indicated). **(f)** Association of *DEAR4* transcript levels with seedling shoot area size (*R* = 0.28, Pearson *p* = 6.7 × 10^-3^, n = 88). **(g)** Association of *FRO6* transcript levels with seedling shoot area size (*R* = 0.62, Pearson *p* = 1.7 × 10^-10^, n = 88). **(h)** Association between seedling size and the expression patterns of the 130 or 29 leaf maturation-induced genes that were eithers induced (orange) or repressed (green) by drought stress.

We found that the severity of HA stress had a dose-dependent effect on leaf maturation gene expression patterns. This was revealed by assessing maturation-induced genes that were induced by drought stress both *(i)* within our HA assay and *(ii)* within our leaf maturation experiment (**Fig. S9**). For example, we identified 130 genes in the mesophyll cell-type that were induced by HA stress in seedlings as well as during leaf maturation, including senescence associated genes such as *RD29A* and *DREB AND EAR MOTIF PROTEIN 4* (DEAR4) (*22, 23*). These genes displayed dose-responsive gene expression patterns in response to HA, with their level of induction over time increasing proportionally to HA stress intensity (**Fig. 4d**). Similar genome-wide expression trends were observed in other cell types (**Fig. S9, Table S7**). Collectively, these data indicate that genes involved in leaf maturation are induced in a dose-responsive manner in response to stress.

We note we also identified dose-responsive expression patterns among smaller set of genes upon which drought stress acted ‘antagonistically’, i.e. maturation-induced genes that were repressed by drought (**Fig. 4e**, **Fig. S9**). For instance, in the mesophyll cell type, 29 genes that were upregulated during leaf maturation but suppressed by drought stress displayed dose-dependent gene expression trends (**Fig. 4e**). Among these genes were those involved in carbon fixation (*CP12 DOMAIN-CONTAINING PROTEIN 1* (CP12)) and sucrose synthesis (*FRUCTOSE-1,6-BISPHOSPHATASE* (FBP)).

### Cell-type specific overexpression of FRO6 promotes leaf growth under stress

*Arabidopsis* adapts to drought stress by decreasing leaf surface area and shoot size (**Fig. 4a**). Lastly, we investigated whether dose-responsive gene expression changes in leaf maturation correlated with changes in shoot size. When examining genes that were induced by leaf maturation and advanced by HA stress, such as *DEAR4*, we discovered that their expression patterns were only weakly associated with changes in seedling size (-0.13 < *R* < 0.54, Pearson) (**Fig. 4f**, **4h**). In contrast, we found that the smaller set of genes that were induced by leaf maturation but repressed by HA stress (i.e. ‘antagonistically’ expressed), such as *FERRIC REDUCTION OXIDASE 6* (FRO6), were more strongly associated with seedling size (0.43 < *R* < 0.89) (**Fig. 4g**, **4h**). This was because the expression profile of antagonistically expressed genes more closely resembled that changes in shoot size (**Fig. 4a**, **4e**). Consequently, while drought’s dominant effect was to advance leaf maturation gene expression, the smaller cohort of maturation induced genes that drought repressed seemed more closely linked to changes in leaf growth in response to stress (i.e., plasticity).

To validate this hypothesis, we aimed to test whether perturbing a gene whose expression was induced during leaf maturation but suppressed by drought would alter leaf growth responses to stress. Among the 29 candidate genes in the mesophyll cell type, we selected FRO6 as a candidate (Fig. 4d), as its expression was most significantly suppressed by drought during leaf maturation (adj. *p* = 2.01 × 10^-19^, linear model) (**Fig. 5a**). FRO6 is a membrane bound ferric chelate reductase that converts iron(III) to iron(II) (*24*), and is hypothesized to transport iron(II) to chloroplasts for energy production (*25*). When we examined subcellular localization of FRO6 using a fluorescent reporter line (*FRO6p*::FRO6-GFP), it appeared to be localized to the endoplasmic reticulum and near chloroplasts within mesophyll cells (**Fig. 5b, Fig. S10**) (*26*).

**Fig. 5:**
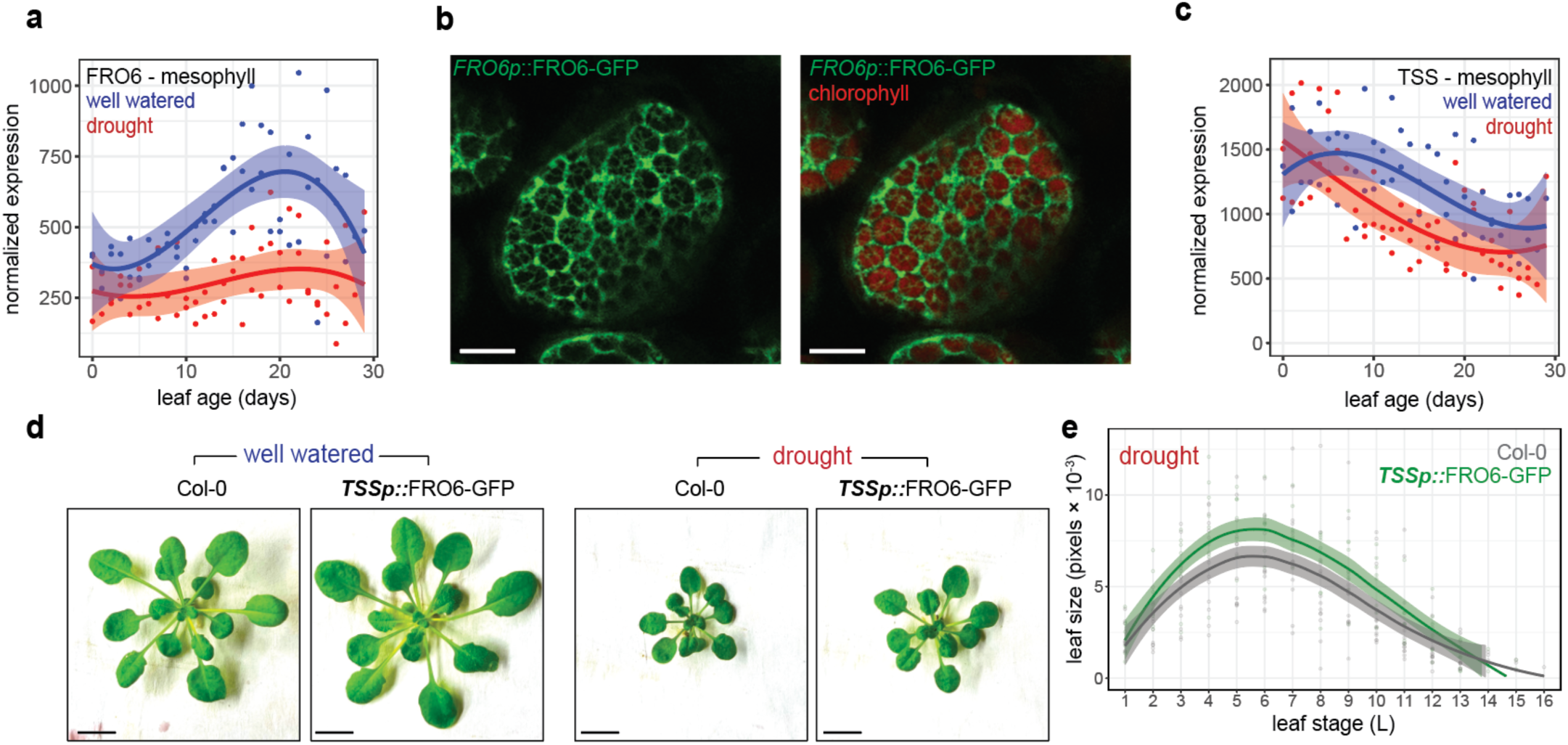
Cell-type specific over-expression of FRO6 within the mesophyll promotes shoot growth under drought stress. **(a)** Mesophyll FRO6 transcriptional abundance under well-watered (100% WC, blue) and drought conditions (22 - 48 % WC, red) across different leaf ages (curve fit using quadratic model, 99% confidence interval indicated). **(b)** Confocal microscopy showing *FRO6::*FRO6-GFP and chlorophyll localization within a mesophyll cell within cotyledons (10μm size indicated). **(c)** Mesophyll TSS transcriptional abundance under well-watered (100% WC) and drought conditions (22 - 48 % WC) across different leaf ages (curve fit using quadratic model, 99% confidence interval indicated). **(d)** Images of Col-0 and *TSSp::*FRO6-GFP rosettes grown under either well-watered or drought conditions. **(e)** Individual leaf area size of Col-0 and *TSSp::*FRO6-GFP vermiculite grown rosettes after 8 days of drought stress (*p* = 6.3 × 10^-3^, ANCOVA model, n = 8 - 15 individuals).

To counter drought-induced transcriptional repression of FRO6 in mesophyll cells, we leveraged our transcriptional atlas to identify gene promoters with high and cell-type specific expression in the mesophyll. Among these, the *TPR-DOMAIN SUPPRESOR OF STIMPY* (*TSS*, also called *REDUCED CHLOROPLAST COVERAGE 2*) (*27*) stood out for its mesophyll-restricted expression that exceeded FRO6 levels by approximately 2-fold (**Fig. 5c, Fig. S10**). We over-expressed FRO6 in a mesophyll specific manner using the 1643 bp upstream region of *TSS*. Notably*, TSS* itself was neither significantly repressed by drought nor by HA stress (adj. p > 0.05, linear model), nor was *TSSp* active within root tissue (**Fig. S10**). Compared to wild-type plants, we observed that two independent transgenic *TSSp::*FRO6-GFP lines grown on vermiculite displayed significantly increased leaf area and shoot biomass under drought conditions (**Fig. 5d, 5e, Fig. S10**). Furthermore, we found that this transgenic line displayed increased shoot area under HA stress compared with wild-type (**Fig. S10**, *p* = 7.3 × 10^-4^, ANCOVA). Collectively, these data indicates that mesophyll-specific over- expression of FRO6 results in enhanced shoot growth under drought conditions in *Arabidopsis*. More broadly, these findings suggest that cell-type specific targeted over-expression of genes induced during leaf maturation but repressed by drought may promote leaf growth under stress.

## Discussion

Upon encountering drought, *Arabidopsis* advances transcriptional responses related to leaf maturation and aging. This finding helps unify earlier reports detailing the impact of drought on leaf development. Elegant studies have shown that drought modifies gene regulation to restrict proliferative and expansive phases, arrest the cell cycle, and hasten the transition to endoreduplication, along with activating senescence-associated genes to expedite leaf death (*1-3, 8, 28*). Here, we demonstrate that these seemingly distinct leaf developmental responses to drought can be understood through the premature induction of transcriptional responses related to leaf maturation, whereby the induced leaf ‘aging’ ultimately limits leaf growth. The advancement of leaf biological age ahead of calendar age has been described elsewhere at the epigenetic level (*29*).

A defining feature of *Arabidopsis* leaf developmental plasticity is its ability to scale in size with stress intensity. The dose-dependent transcriptional responses to stress we describe here are associated with this plasticity. Examples of dose-responsive transcriptional changes related to organ size are observed elsewhere, such as in response to nutrient availability in *Arabidopsis* roots (*30*), and water stress responses in *Arabidopsis* seedlings and rice shoots (*31, 32*). We hypothesize that such dose-responsive gene expression may, in part, depend on cellular hormone levels. Specifically, we find that genes responsive to cytokinin and salicylic acid treatment - hormones whose levels are known to prolong or extend leaf lifespan, respectively (*15, 18*) - are well represented among leaf maturation genes that change in response to drought stress.

Our findings carry important implications for developing drought resilient crops. For instance, substantial efforts have focused on engineering crop varieties that can withstand drought stress while maintaining optimal leaf growth under non-stress conditions (*33*). This has proven to be challenging, as candidate genes identified through drought response assays often improve drought resilience at the expense of overall plant stature (*34, 35*). Here, we show that a significant component of drought responsive gene expression indeed works to advance leaf maturation to reduce leaf size. Surprisingly, the less common maturation gene expression responses that are antagonistically regulated by drought stress, such as those exhibited by FRO6, may represent more promising candidates for developing traits that can withstand drought stress without negatively impacting shoot growth. Our transcriptional atlas of leaf development may facilitate similar cell-type specific strategies in the future.

## Materials and Methods

### *Arabidopsis* growth conditions for rosette leaf size quantification and single-nuclei transcriptome sequencing

*Arabidopsis* Col-0 seeds were surface sterilized and stratified for 2 days, and then grown for 17 days on LS media (Cassion). Plates were supplemented with 1% sucrose, and incubated under short day conditions (8h light) with light intensity set at 150 μmoles at 22°C. After this period, plants were transferred to vermiculite and grown on 0.75× LS media without sucrose for 12 days, maintaining constant saturation. On the 12th day, the first leaf samples was taken for sci-RNA-seq3 sequencing, and collected 4.5 hours after subjective dawn. To introduce a drought stress, excess water was removed until all pots reached field capacity (FC). We note that in the context of this manuscript, 100% FC is synonymous with 100% water content (WC). Subsequent leaf sampling was conducted daily for 8 days at the same time of day, and degree of evaporation measured by weighing each pot. Three rosettes were sourced from drought-stressed and three from well-watered control groups per time point. 2 biological replicates were performed (i.e. two distinct drought stress time course experiments performed on two separate occasions). At the time of sampling, whole rosettes were flash-frozen, before each leaf was excised and placed into a 96-well plate. Leaf measurements were conducted using Plant Growth Tracker software (https://github.com/jiayinghsu/plant-growth-tracker/tree/main/cascade_rcnn) (**Table S4**). We applied an ANCOVA model (designating drought stress and time as qualitative and quantitative variables respectively) to assess statistical differences between rosette shoot size. We note that *TSSp::*FRO6- GFP genotypes were grown under the same vermiculite growth conditions as described above.

### *Arabidopsis* growth conditions for exogenous hormone treatment

*Arabidopsis* seedlings were sterilized and stratified as described above before being grown on vertical plates for 17 days under short day conditions on 1× LS media supplemented with 1% sucrose. After this time, seedlings were transferred to vermiculite (0.75× LS media, no sucrose). Plants were grown at complete FC saturation for 18 days. On the 18^th^ day, 2 hours after subjective dawn, both roots (through replacing growth media) and shoots (through foliar spray) were treated with one of the following hormone treatments: 10 µM (±)-Abscisic acid, 25 µM 2,4-Dichlorophenoxyacetic acid, 50 µM Gibberellin A_3_, 300 µM Salicylic acid, 1 µM *trans*-Zeatin, 10 nM Brassinollide, 10 µM Methyl jasmonate, 50 µM 1-Aminocyclopropane-1-carboxylic acid (ACC). Each hormone solution contained 0.01 % DMSO and 0.1 % ethanol. A mock treatment control was also included. Plants were treated for ∼3 h, before rosettes flash frozen in liquid nitrogen. ∼ 15 plants were collected per treatment.

### *Arabidopsis* growth conditions for ‘Hard Agar’ (HA) treatment

*Arabidopsis* Col-0 seeds were sterilized and stratified as described above before being grown on vertical plates supplemented with 1 × LS media, 1% sucrose and 2% agar with a light period of 8 hours (150 μmoles) at 22°C. We designated 2% agar and 1 × LS media as “1×HA” dose. Thus, a “2×HA” dose consisted of 4% agar and 2×LS media. Additional description and validation of HA stress is described in Gonzalez *et al.* (*21*). Sampling of individual seedlings for sci-RNA-seq3 sequencing commenced on the 15th day, 4.5 hours after subjective dawn, and continued for a total of 11 days. Plants were imaged on plates for shoot area measurements, before excising roots away, and flash freezing each individual shoot in a 96-well plate (n = 7 individual seedlings per time point, per HA dose). Images were processed using Plant Growth Tracker software (**Table S4**). We note that for *TSSp*::FRO6 shoot area measurements, the same HA growth protocol was followed as described above however images were taken 29 days after sowing (n= 6 individual seedlings across two plates, per treatment, per genotype).

### Nuclei extraction and single nuclei RNA sequencing (sci-RNA-seq3)

sci-RNA-seq3 was performed as described in Cao, *et al*. (*9*) with the following notable exceptions. Each frozen leaf or seedling sample was bead bashed (Qiagen) in a 96 well plate format. The resulting frozen homogenate was resuspended in resuspension buffer (10 mM Tris-HCl pH 7.4, 10 mM NaCl, 3 mM MgCl_2_, 1% PBS, 0.5% DEPC). Tissue samples were then passed through a 96-well 30-μm filter. Washed nuclei were concentrated and nuclear RNA reverse-transcribed with a well specific primer. Subsequent ligation, tagmentation and PCR steps of sci-RNA-seq3 were followed as described in Cao, *et al*. Libraries were sequenced on the Illumina Novaseq 6000 with 150 bp paired-end chemistry. Resulting reads were aligned to the Arabidopsis TAIR10 genome with Araport11 annotation (*36*). Number of nuclei sequenced per sample, and UMI per nuclei are reported in **Table S1.**

### Nuclei extraction and single nuclei RNA sequencing (10X RNA-seq)

Nuclei isolation was performed upon whole rosettes. Frozen tissue was crushed using a mortar and pestle, and nuclei released from homogenate using a resuspension buffer (10 mM Tris-HCl pH 7.4, 10 mM NaCl, 3 mM MgCl_2_, 1% PBS, 1% Superase RNAse Inhibitor). The resulting homogenate was filtered using a 30μm filter. To enrich for nuclei, an Optiprep (Sigma) gradient was employed. Enriched nuclei were then purified using Fluorescent Activated Cell Sorting (FACS). Purified nuclei were loaded directly onto the 10X machine 10X-Gene Expression v3.0 chemistry and sequenced on the Illumina Novaseq 6000 with 150 bp paired end chemistry. Libraries were sequenced aligned to the Arabidopsis TAIR10 genome with Araport11 annotation (*36*). Chloroplast and mitochondrial reads were removed. Number of nuclei per sample, and UMI per nuclei are reported in **Table S1.**

### Nuclei clustering

Transcriptional atlases from each experiment were assembled using Seurat (*37*). Nuclei were first subsetted using a minimum UMI threshold of 450 reads. Then, nuclei from across different experiments (i.e. from individual rosette leaves, individual seedlings, and hormone treatments) were combined into one atlas. The integrated dataset was subjected to clustering, employing the top 3000 variable features that were shared across all datasets. Subsequent UMAP projections were constructed using the first 30 principal components. Very small clusters were considered artifact and removed. Cell types were subsequently annotated using marker genes listed in **Table S2**. The exception was the bundle sheath cell type cluster, where we overlapped the bundle sheath specific markers described in Kim *et. al* (*38*) (enrichment score > 6) with the cell makers present in our atlas (enrichment score >3). We note that UMAP projections of cell types under well-watered conditions (100% water content) or drought conditions (< 39% water content) were performed using the same approach as described above with the notable exception that we down sampled nuclei to ensure equal numbers of nuclei were present across well-watered and drought samples of nuclei within each leaf stage, and used genes found differentially expressed during leaf development or responsive to drought stress as variable features.

### Detecting cell-type specific leaf maturation gene expression patterns, and their response to stress

To detect genome-wide changes in gene expression during leaf maturation, we established two criteria. First, a gene needed to be differentially expressed across different leaf stages within a rosette. Second, to exclude genes associated with leaf morphology alone (i.e. heteroblasty), a gene also needed to be differentially expressed over time as each leaf matured. We used a linear model to test these criteria. Initially, we examined gene expression patterns in whole leaf organs using a multivariate linear model:

*gene expression_a_* = leaf stage + time + drought + *c*

This analysis, which included data from both well-watered and drought-stressed conditions, indicated that the factors ‘leaf stage’ and ‘time’ were co-linear. Thus, we combined these two variables into a single ‘leaf age’ variable, defined by the following formula:

*leaf age* = 23 + time + (leaf stage / 2) + *c*

Following this, we modeled gene expression profiles for each cell type using a simplified model:

*gene expression_a_* = leaf age + drought + *c*

Where both time and leaf stage were quantitative variables (see **Fig. S4**). This multivariate linear model was implemented in DESeq2 (*39*) on each cell-type’s pseudo-bulked gene expression profile, where genes were first quantile normalized. We classified a gene as involved in maturation if the leaf age coefficient was significant (adjusted *p* < 0.01, leaf age coefficient > 0.04), and as drought-responsive if the drought coefficient was significant (adjusted p < 0.01).

### Detecting hormone responsive genes among cell-type classes

Genes found differentially expressed in response to exogenous hormone treatment were detected by comparing each hormone treatment to the mock control, using the FindMarkers() command in Seruat (*37*) (adj. *p* <0.1, likelihood-ratio test). Because less abundant cell types did not have enough nuclei to perform robust statistical testing, we combined vasculature cell types together into a ‘cell class’ (i.e. phloem, xylem, bundle sheath, hydathode and myrosin idoblast cell types). We used the same approach to combine epidermal cell types together (i.e. epidermal, guard and trichome cell types). In each cell class we found some genes that were differentially expressed in response to multiple hormone treatments. To ensure we only analyzed genes that were responsive to specific hormones, for each cell class we removed genes that were differentially expressed in response to 3 or more hormones. We intersected our final set of hormone responsive genes with those that were found differentially expressed during leaf maturation or in response to drought within our individual leaf rosette experiment (linear model, adj. *p* <0.01). We only retained genes whose intersect was significant for further analysis (Fisher exact test *p* < 0.05, using a background of all expressed genes). To ensure greater stringency of statistical testing, intersections were performed with directionality. Gene Ontology (GO) terms were called using agriGO software, using the whole genome as background.

### Detecting dose-responsive cell-type specific transcriptional responses to HA stress

We used a multivariate linear model to identify genes that were both differentially expressed during the 11 days of *Arabidopsis* seedling shoot growth, as well as dose-response to the level of HA stress. This was achieved by first pseudo-bulking each cell-type’s expression profile across each of the 88 conditions assayed (11 time points, 8 HA doses), and removing low count reads. Then, the statistical model was implemented in DESeq2 (*39*) using quantile normalized reads, and where time (days) and HA dose were considered quantitative variables. A gene was identified as significantly differentially expressed to both factors at an adjusted p-value threshold of 0.01. This modeling approach was implemented for 6 of the major leaf cell types identified, where resulting lists of significant genes are found in **Table S7.** We binned genes as either HA stress induced or HA stress repressed by relying on whether the HA factor’s coefficient was positive or negative respectively. To assess agreement between *(i)* the gene expression responses elicited by HA stress and *(ii)* leaf maturation gene expression responses changes to drought stress we performed a Fisher exact test across these two datasets in a cell-type specific manner, using all expressed genes within the atlas as background.

### Plasmid construction

A 4198 bp genomic fragment from the FRO6 locus, which contains the FRO6 coding region, 886 bp upstream of ATG, and 211 bp of 3’ UTR, was used to generate *FRO6p*::FRO6:GFP transgenic line. GFP was translationally fused to the C-terminus of FRO6 before the stop codon. For the *TSSp*::FRO6:GFP transgenic line, a 1643 bp fragment upstream of TSS start codon was used to replace the FRO6 promoter. The same TSSp fragment was used in generating the *TSSp*::GUS transcriptional fusion. All transgenes were cloned into the binary vector pMX202 and stably transformed into *Arabidopsis* Col-0 background.

### Confocal Microscopy

Cotyledons of 6-day-old seedlings carrying the *FRO6*::GFP fusions were imaged with a Leica Stellaris 8 confocal microscope, using 473 nm laser excitation. GFP fluorescence signals were collected between 480-580 nm, and the chlorophyll autofluorescence in this range was excluded using the tau-gating function. The red chlorophyll autofluorescence was collected between 600-670 nm.

### GUS Staining

GUS activity staining was carried out as described in (*40*), using 10 mM potassium ferro and ferri cyanide. The GUS-stained seedlings were mounted in 30 % glycerol and imaged using a Zeiss Axio Zoom.V16 stereomicroscope equipped with an axiocam 305 color camera.

## Data and Code Availability

Single-cell RNA-seq data are publicly available through GEO (GSE290214). Nuclei data are hosted at the Klarman Cell Observatory at: https://singlecell.broadinstitute.org/single_cell/study/SCP2703). Microscopy data reported in this paper will be shared by the lead contact upon request.

## Author Contributions

J.S., J.R.E., X.W. and J.C designed the experimental plan. J.S. performed the nuclei sequencing experiments and genomic analyses. X.W. performed transgenics and microscopy. J.S., J.X. and N.I.E. performed physiological measurements. T.J. assisted in bioinformatics. J.N. performed sequencing workflows. J.S. and J.R.E. wrote the manuscript.

## Supplementary Materials

**Figure S1:** Identifying *Arabidopsis* leaf cell types from sci-RNA-seq3 and 10X experiments.

**Figure S2:** The *Arabidopsis* leaf atlas is hosted by the Klarman Cell Observatory.

**Figure S3:** Cell-type specific transcriptional responses underlying *Arabidopsis* leaf maturation.

**Figure S4:** Associating gene expression responses with leaf maturation.

**Figure S5:** Inducing drought stress and measuring its effect on leaf maturation

**Figure S6:** Drought stress changes leaf maturation transcriptional dynamics.

**Figure S7:** Cell-class specific transcriptional responses to exogenous hormone treatment.

**Figure S8:** Overlapping hormone responsive genes with those differentially expressed during leaf maturation or in response to drought stress.

**Figure S9:** Dose-responsive transcriptional responses to hard agar (HA) stress across 6 major cell types.

**Figure S10:** Targeted upregulation of FRO6 using the *TSSp* mesophyll-specific promoter.

## Supplementary Tables

**Table S1:** Distribution and count of nuclei across leaf and seedlings assayed by sci-RNA-seq3.

**Table S2:** List of validated cell type markers used for cluster annotation.

**Table S3:** Lists of genes differentially expressed during leaf maturation.

**Table S4:** Rosette and seedling physiological measurements.

**Table S5:** Lists of leaf maturation responsive genes differentially expressed in response to drought stress.

**Table S6:** Cell class specific hormone responsive gene lists.

**Table S7:** List of growth-responsive genes differentially expressed in response to HA stress.

---

**Fig. S1:**
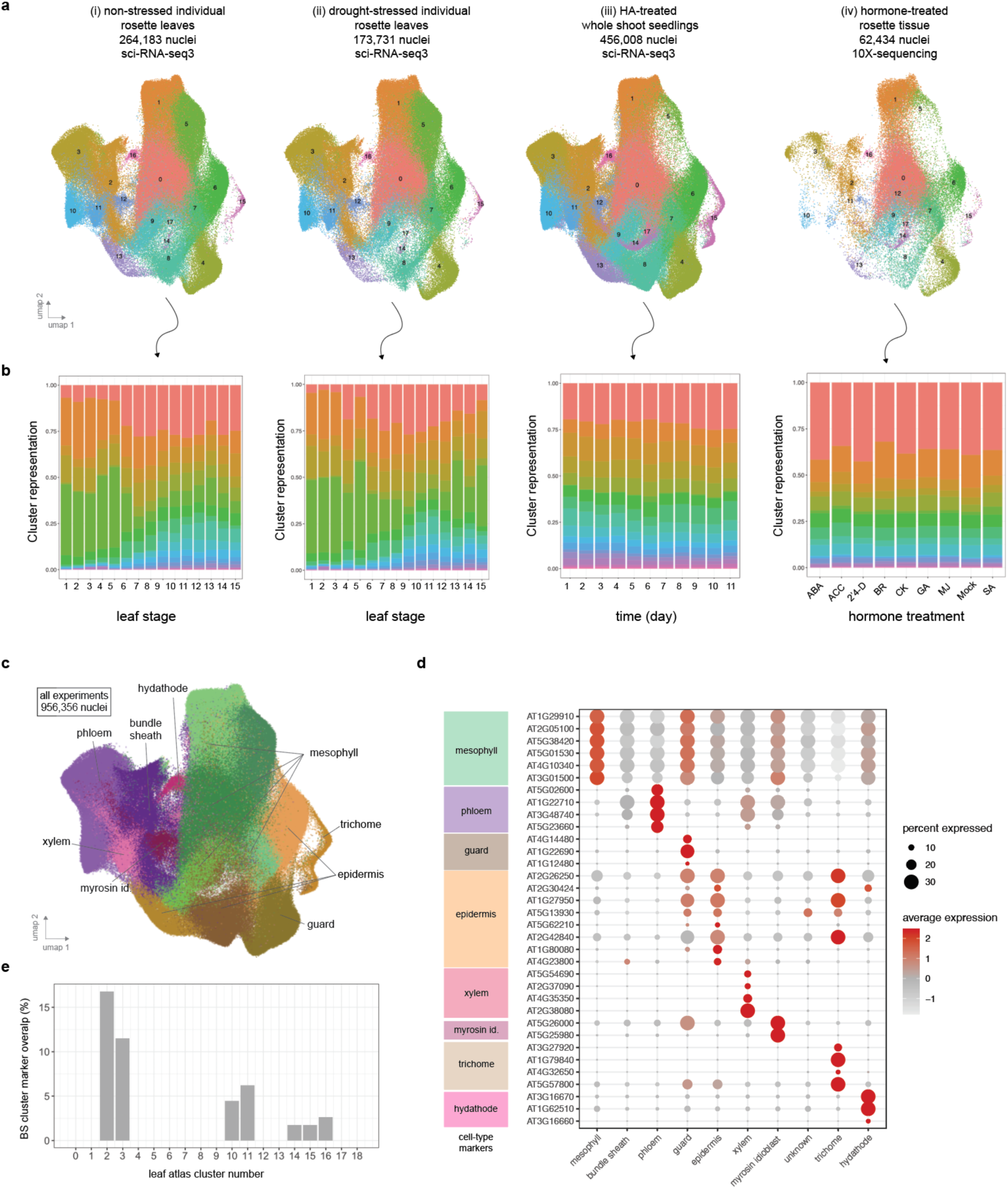
An *Arabidopsis* leaf transcriptional atlas. **(a)** A total of 956,356 nuclei were sequenced across 4 different experiments: (*i*) non-stressed individual rosette leaves, (*ii*) drought-stressed individual rosette leaves, (*iii*) whole shoot seedlings grown on HA agar, and (*iv*) whole rosette tissue treated exogenously with hormones. All four experiments were integrated together into a single atlas before identifying *Arabidopsis* leaf cell types. **(b)** Cluster representation for each leaf stage (rosette experiments), time of sampling (seedling experiment), or hormone treatment. **(c)** Cell type annotation for each cluster within the atlas. **(d)** Cell types were annotated by relying on expression of validated cell type marker genes (see **Table S2**). **(e)** The bundle sheath cell type (cluster 2) was identified by overlapping the significant bundle sheath markers found in *Kim et. al* (*38*) with the cluster- specific marker genes from this study.

**Fig. S2:**
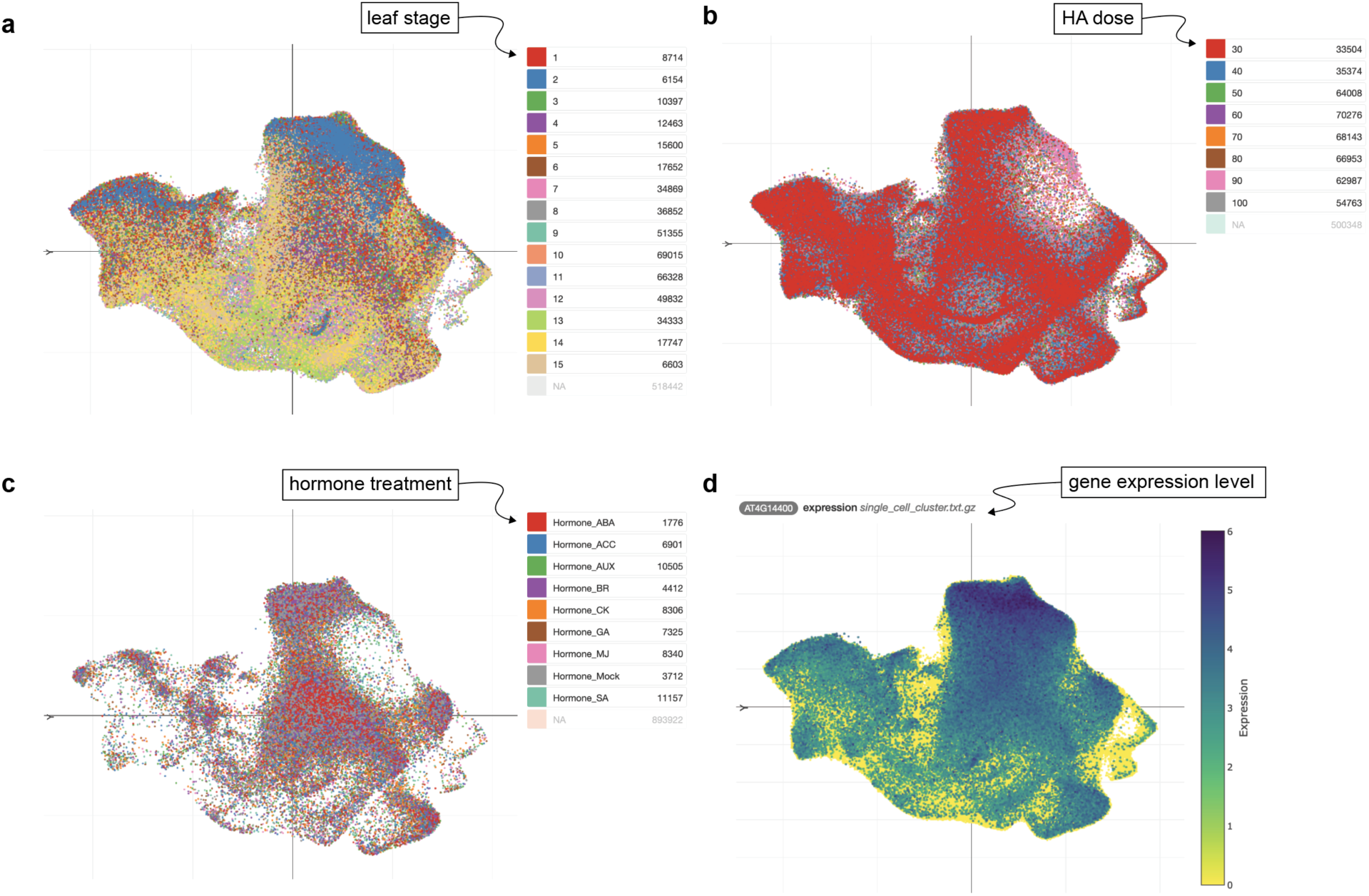
The *Arabidopsis* leaf atlas is hosted online by the Klarman Cell Observatory. 956,356 nuclei spanning the different experiments presented in this study are available online. This interface allows users to view how nuclei cluster by **(a)** different leaf stages, **(b)** growth on different HA doses, **(c)** hormone treatment, as well as **(d)** visualize individual gene expression (ACD6 expression is shown). Additional analyses, such as gene correlation analysis, can be initiated by the user. These data can be accessed at: https://singlecell.broadinstitute.org/single_cell/study/SCP2703.

**Fig. S3:**
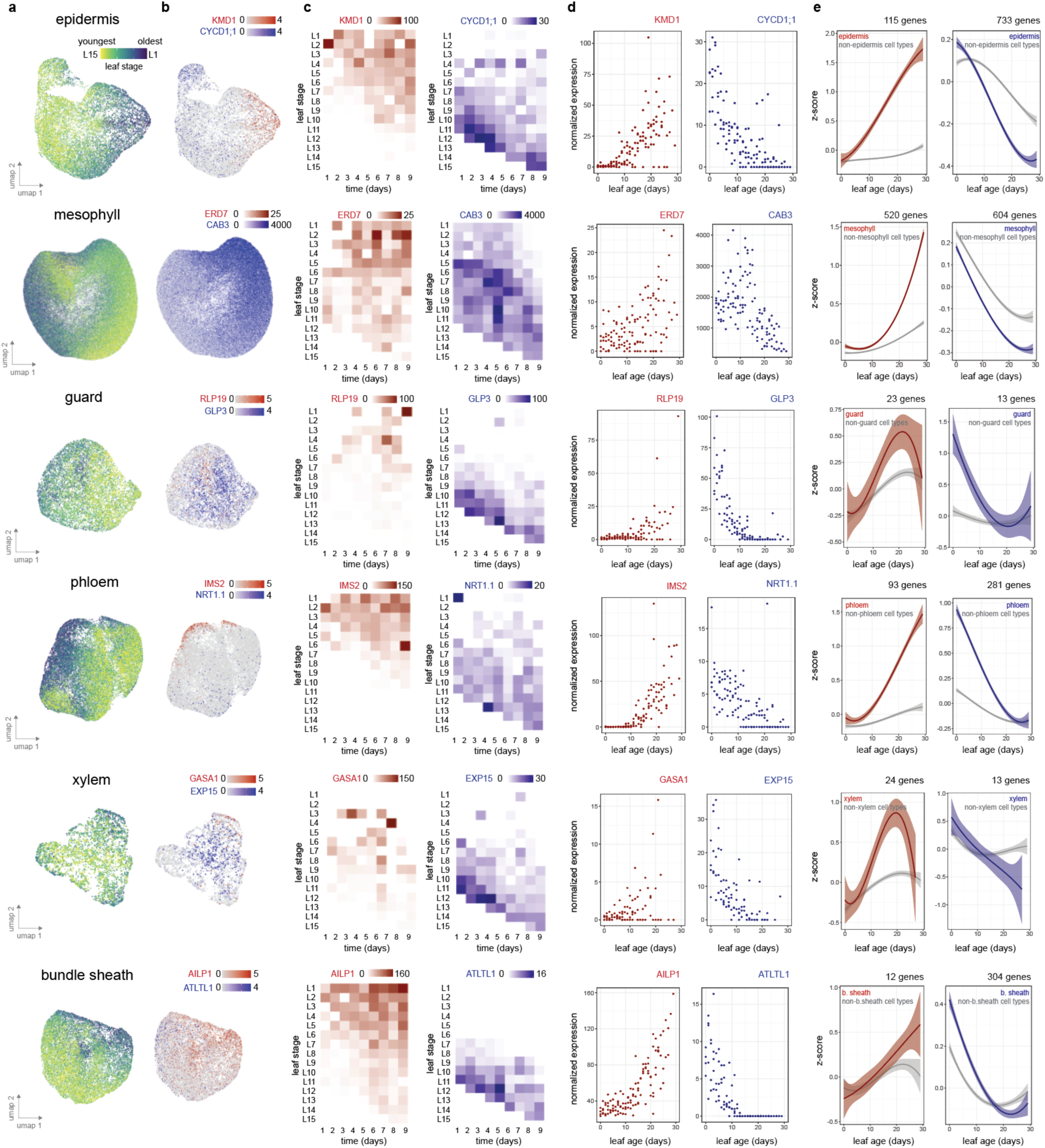
Cell-type specific transcriptional responses underlie *Arabidopsis* leaf maturation. **(a)** Sub- clustering of 6 leaf cell types, with nuclei colored by the leaf maturation stage they originate from. **(b)** Expression profile of selected genes that were either upregulated (red) or downregulated (blue) during leaf maturation within each cell type. **(c)** Cell-type specific pseudo-bulked expression profile of selected genes ordered by leaf stage and day of sampling. **(d)** Expression profile of selected genes as leaves aged and matured. **(e)** z-score normalized expression trend of cell-type specific genes either significantly induced or repressed during leaf maturation (adj. *p* < 0.01, linear model statistic, curve fit using quadratic model, 99% confidence interval indicated).

**Fig. S4:**
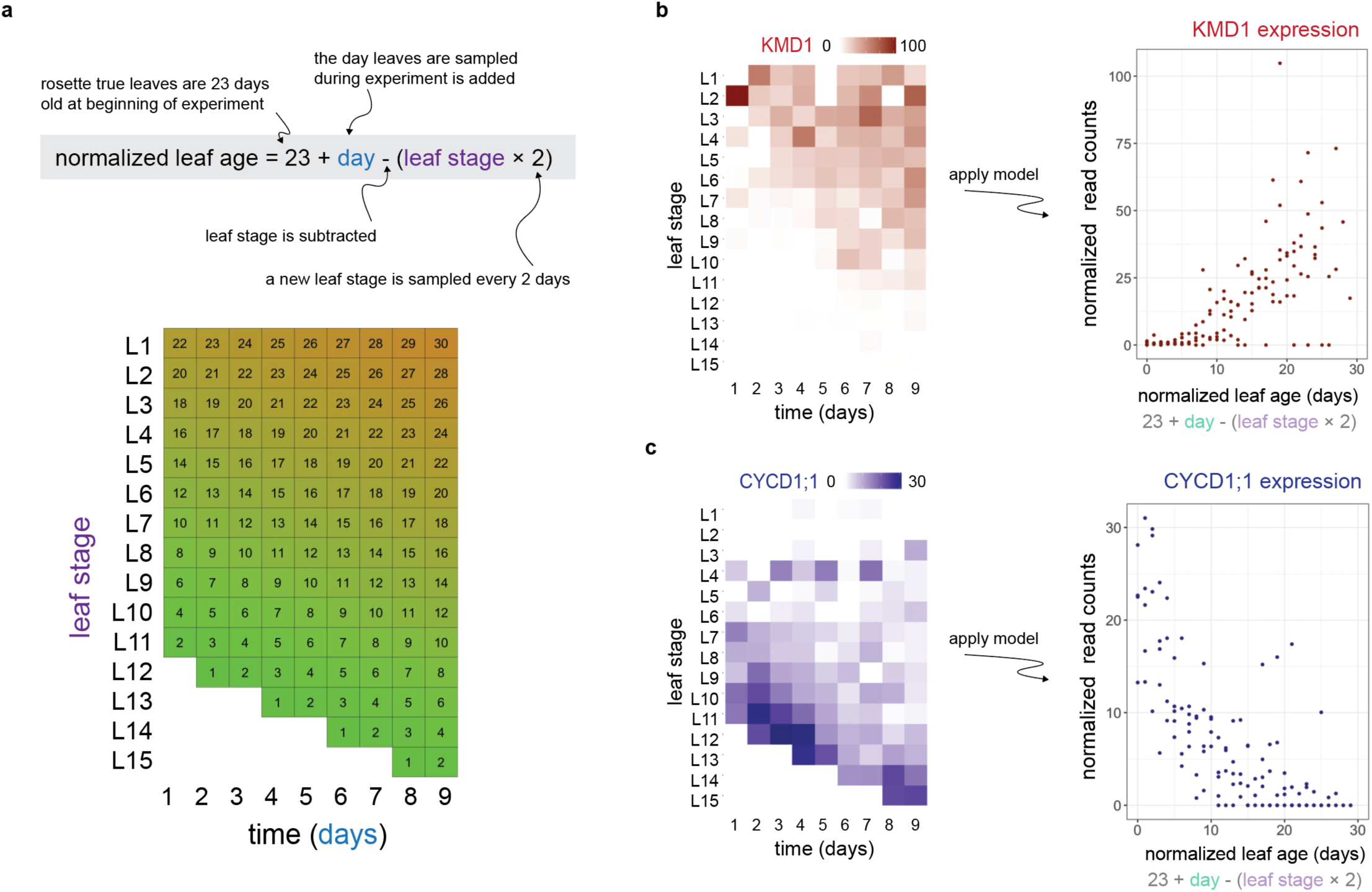
Computing a normalized leaf age value based off a leaf’s developmental stage sampling time. **(a)** Normalized leaf age was calculated by taking into account the age of the rosette leaves at the start of the experiment, the sampled timepoint, and the stage of the sampled leaf. The application of this model to each leaf sample is shown below, where each grid number represents a leaf’s computed age value. This model is applied to the expression patterns of two example genes **(b)** *KMD1* and **(c)** *CYCD1;1*.

**Fig. S5:**
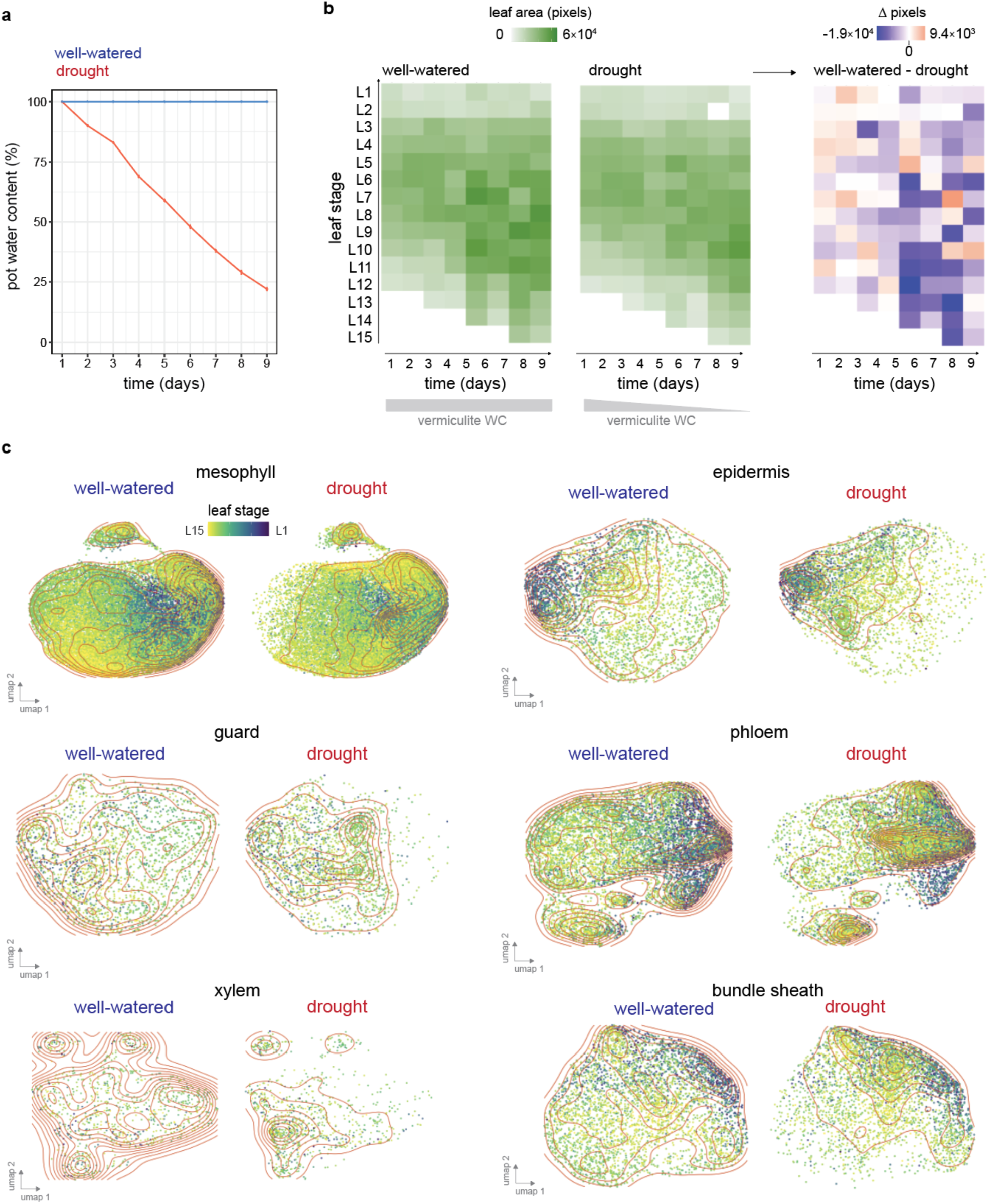
Inducing drought stress and measuring its effect on leaf maturation. **(a)** Pot vermiculite water content measurements of *Arabidopsis* rosettes grown under constant saturated conditions (well-watered), or with water withheld for a 9-day period (drought, n = 12 - 22, standard error indicated). **(b)** Individual leaf area size of rosettes grown under vermiculite conditions. Drought stress had a significant impact on leaf area (3-way ANCOVA *p* = 1 × 10^-3^, n = 1 - 3). The difference in leaf size (τι pixels) is displayed on the right, where blue and red indicate a reduction or increase in leaf size respectively **(c)** Equal numbers of nuclei from 6 cell-types sourced from well-watered or drought conditions subclustered and colored by their respective leaf stage (red contour lines represent levels of equal point density).

**Fig. S6:**
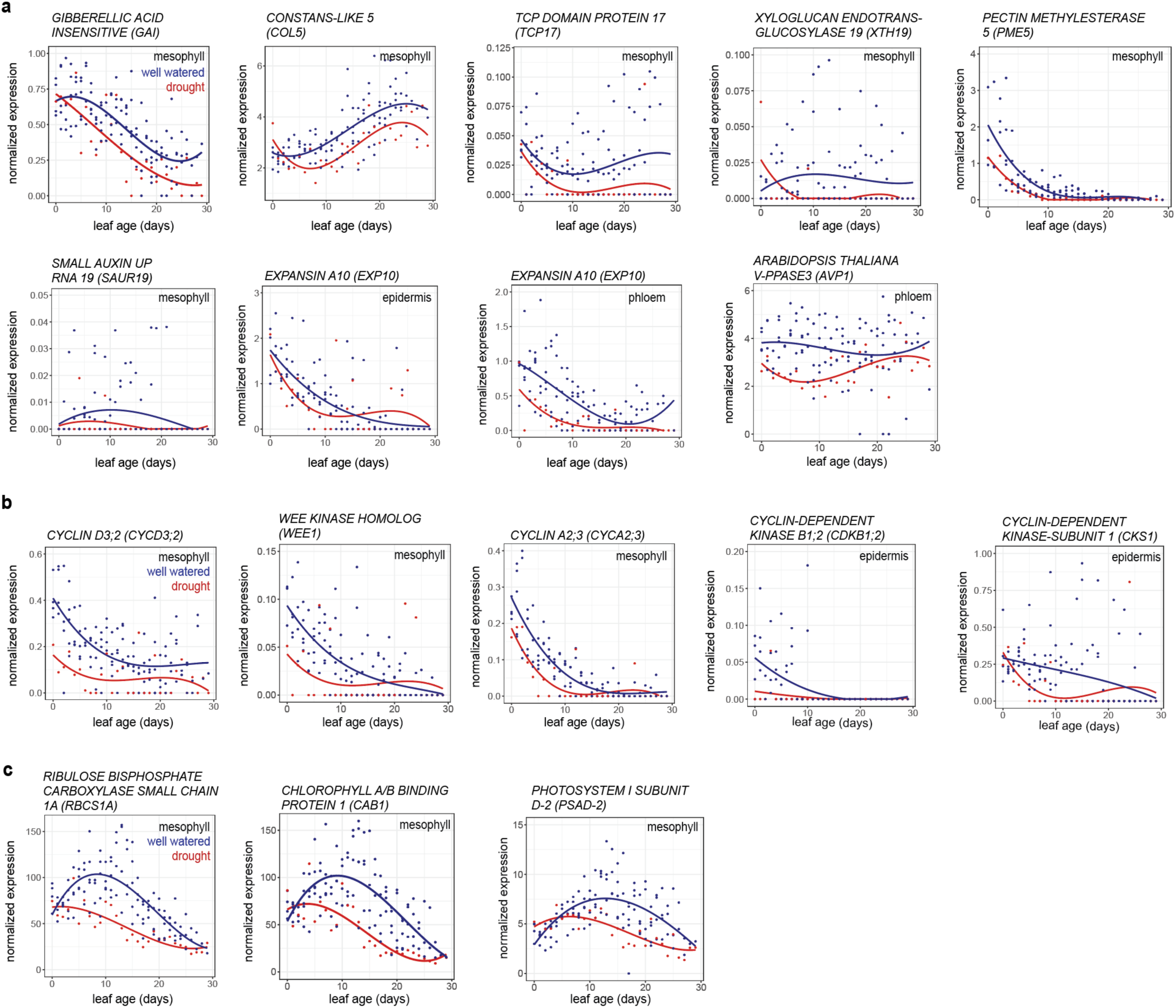
Drought stress changes leaf maturation transcriptional dynamics. Cell type specific expression patterns of genes involved in either **(a)** leaf development, **(b)** cell cycle, **(c)** photosynthesis, under either well- watered or drought conditions (curve fit using quadratic model).

**Fig. S7:**
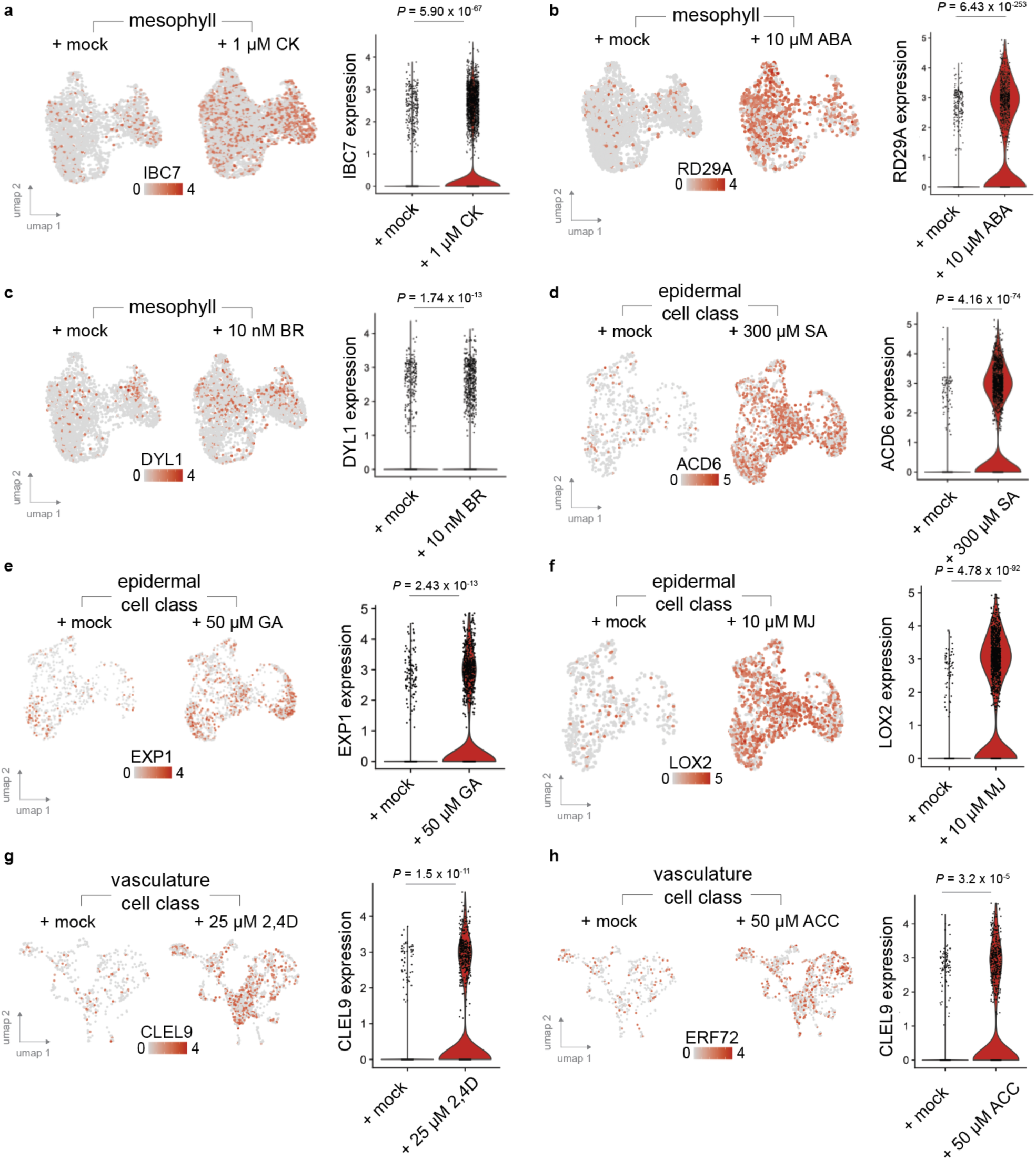
Cell-class specific transcriptional responses to exogenous hormone treatment. UMAPs and accompanying violin plots display a candidate gene’s differential expression patterns in response to a hormone treatment within a cell-type class. **(a - c)** The mesophyll cell-type differentially expresses either *INDUCED BY CYTOKININ 7* (IBC7), *RESPONSE TO DESICCATION 29A* (*RD29A*), or *DORMANCY-ASSOCIATED PROTEIN-LIKE 1* (DYL1) in response to *trans*-Zeatin (CK), abscisic acid (ABA), or brassinollide (BR), treatment respectively. **(d - f)** The epidermal cell-class (combining epidermal, guard and trichome cell types) differentially expresses either *ACCELERATED CELL DEATH 6* (ACD6), *EXPANSIN A1* (EXP1), or *LIPOXYGENASE 2* (LOX2) in response to salicylic acid (SA), gibberellin (GA) or methyl jasmonate (MJ), treatment respectively (likelihood ratio test). **(g - h)** The vasculature cell-class (combining phloem, xylem, bundle sheath, hydathode, myrosin idioblast cell types) differentially express either *CLE LIKE 9* (CLEL9) or *ETHYLENE RESPONSE FACTOR 72* (ERF72) in response to either synthetic auxin (2,4-D) or ethylene precursor 1-Aminocyclopropane- 1-carboxylic acid (ACC) respectively. Additional hormone responsive genes are listed in **Table S6**.

**Fig. S8:**
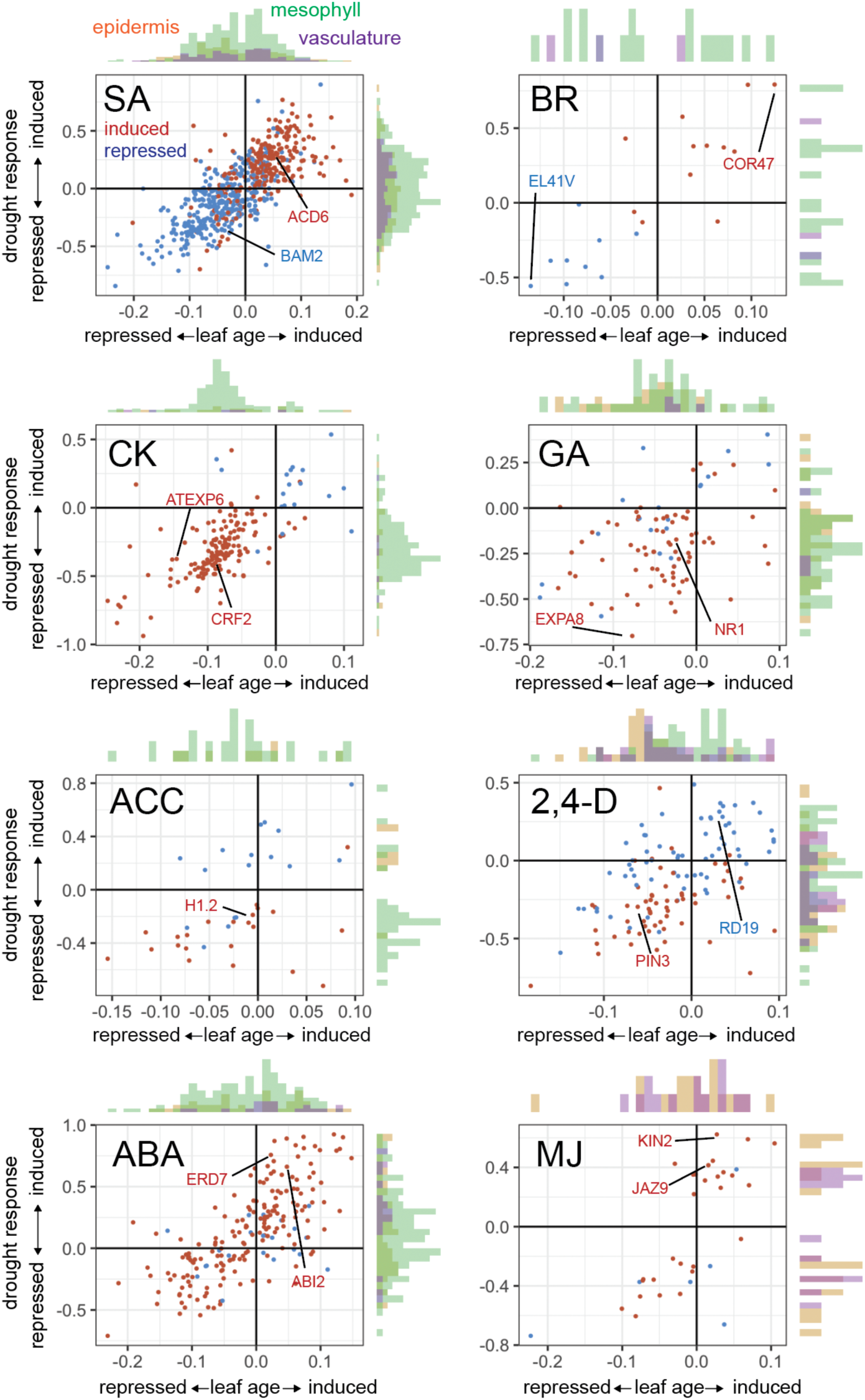
Overlapping hormone responsive genes with those found differentially expressed during leaf maturation or in response to drought stress. Induction (red) or repression (blue) of genes significantly differentially expressed in response to hormone treatment (adj. *p* < 0.1, likelihood ratio test), and each gene’s respective induction or repression in response to leaf maturation or drought stress (adj. *p* < 0.01, linear model, axis units are coefficients of linear model). Histograms indicate in which cell-type class the hormone response was detected. Salicylic acid (SA), brassinosteroid (BR), cytokinin (CK), gibberellin (GA), ethylene precursor 1- aminocyclopropane-1-carboxylic acid (ACC), synthetic auxin (2,4-D), abscisic acid (ABA), methyl jasmonate (MJ). Names of example genes responsive to each hormone indicated (additional genes are listed in **Table S6**).

**Fig. S9:**
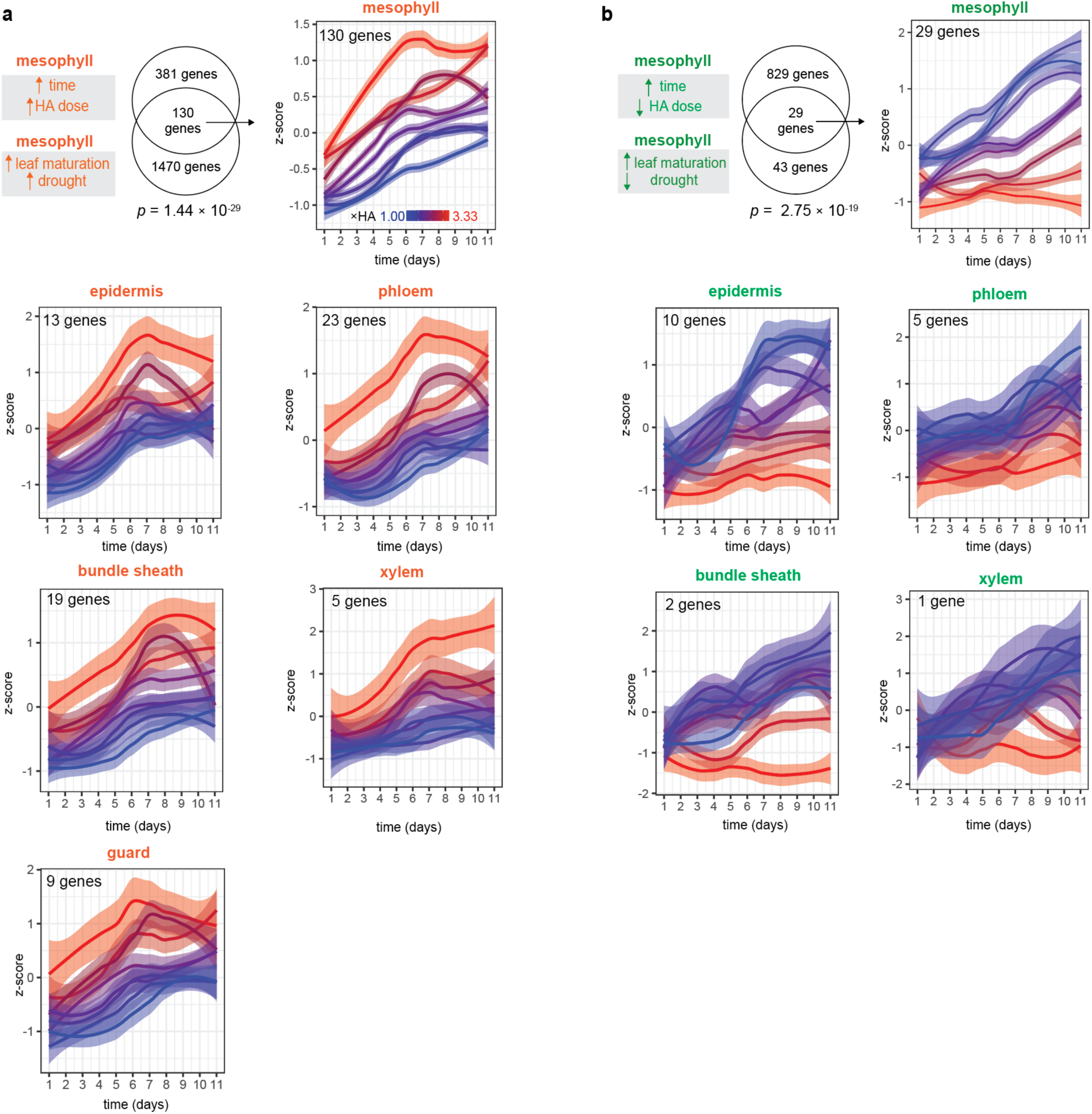
Dose-responsive transcriptional responses to hard agar (HA) stress across 6 cell types. **(a)** Overlap of genes in each cell type that were induced during development and induced by drought stress both *(i)* within the HA time course assay and *(ii)* within the leaf maturation experiment. The significance of this overlap was assessed using a Fishers Exact Test (for conciseness, only mesophyll intersect is displayed). Gene expression responses within this intersect are presented as z-score plots (line fit using locally estimated scatterplot smoothing, 99% confidence interval indicated). **(b)** Overlap of genes in each cell type that were induced during maturation and repressed by drought stress both *(i)* within the HA time course assay and *(ii)* within the leaf maturation experiment.

**Fig. S10:**
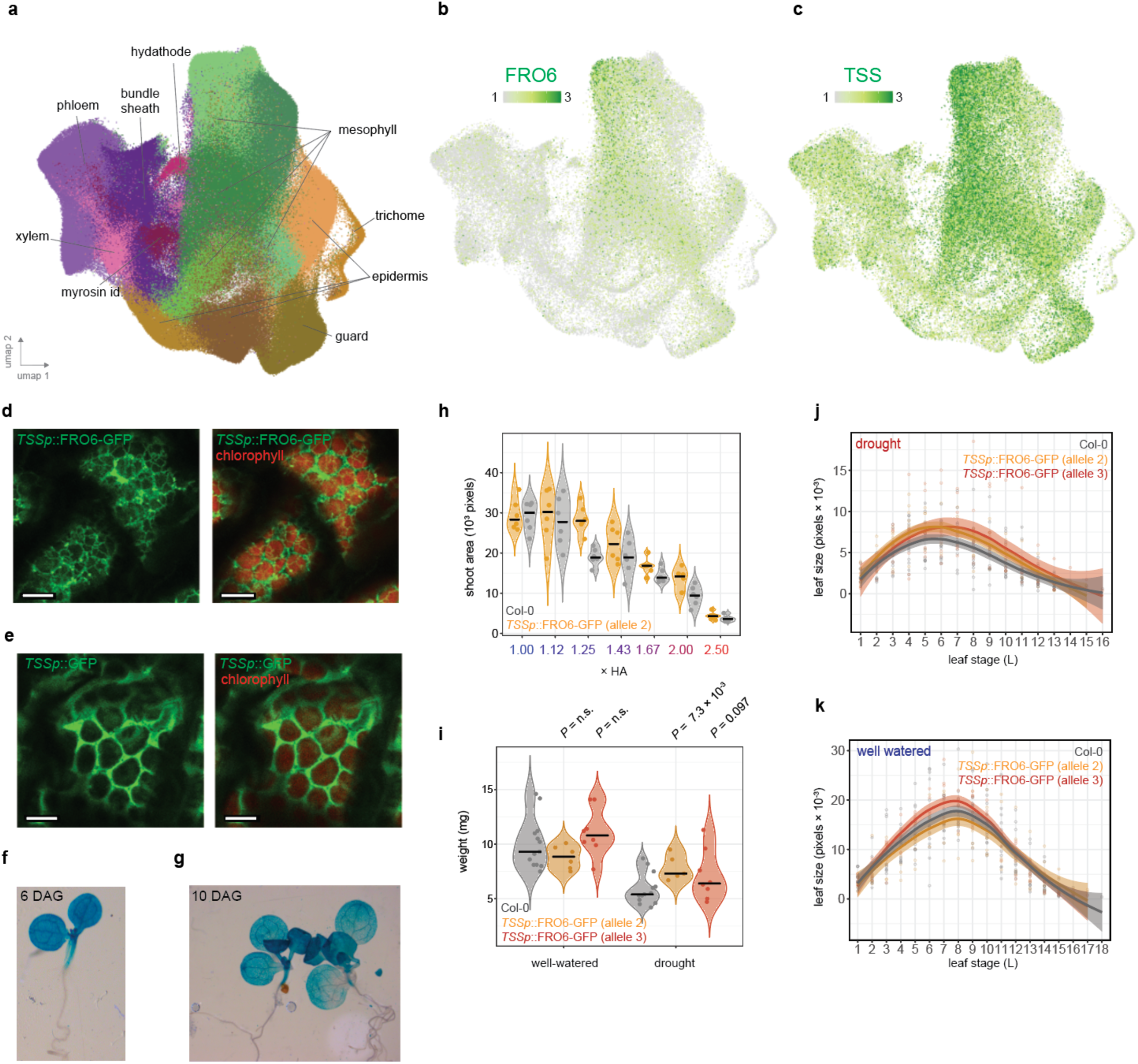
Cell-type specific upregulation of FRO6 using the *TSSp* mesophyll-specific promoter. **(a)** Cell type annotation of each cluster within the atlas. Expression pattern of **(b)** FRO6 and **(c)** TSS within the atlas. **(d)** Confocal microscopy images showing *TSSp::*FRO6-GFP (green) and chlorophyll (red) subcellular localization within mesophyll cells (bar represents 10µm). **(e)** Confocal microscopy images of *TSSp*::GFP subcellular localization within mesophyll cells (bar represents 10µm). GUS reporter line of *TSSp* activity in *Arabidopsis* seedlings **(f)** 6 and **(g)** 10 days after sowing (DAS). **(h)** Shoot area of Col-0 and *TSSp::*FRO6-GFP seedlings were grown across increasing stress HA intensity differ significantly (*p* = 7.3 × 10^-4^, ANCOVA model, genotype factor, n = 6 individuals, bar indicates median). **(i)** Whole rosette dry weight of Col-0 and 2 different *TSSp::*FRO6- GFP alleles grown under either well-watered or drought-stress conditions (Welch’s one-sided t-test *p* indicated, n = 5 - 13 individuals, bar indicates median). **(j)** Individual leaf area size of Col-0 and *TSSp::*FRO6-GFP vermiculite grown rosettes after 8 days of drought stress (allele 3 *p* = 6.31 × 10^-3^, allele 2 *p* = 6.34 × 10^-3^, ANCOVA model, n = 8 - 15 individuals). **(k)** Individual leaf area size of Col-0 and *TSSp::*FRO6-GFP vermiculite grown rosettes under well-watered conditions (allele 3 *p* = 0.58, allele 2 *p* = 0.33, ANCOVA model, n = 8 - 15 individuals).

## References and Notes

1. S. Munne-Bosch, L. Alegre, Die and let live: leaf senescence contributes to plant survival under drought stress. Funct Plant Biol 31, 203–216 (2004).

2. A. Skirycz et al., Developmental stage specificity and the role of mitochondrial metabolism in the response of Arabidopsis leaves to prolonged mild osmotic stress. Plant Physiol 152, 226–244 (2010).

3. P. Clauw et al., Leaf Growth Response to Mild Drought: Natural Variation in Arabidopsis Sheds Light on Trait Architecture. Plant Cell 28, 2417–2434 (2016).

4. H. Claeys, D. Inze, The Agony of Choice: How Plants Balance Growth and Survival under Water- Limiting Conditions. Plant Physiology 162, 1768–1779 (2013).

5. Z. He, S. Webster, S. Y. He, Growth-defense trade-offs in plants. Curr Biol 32, R634–R639 (2022).

6. I. Efroni, E. Blum, A. Goldshmidt, Y. Eshed, A protracted and dynamic maturation schedule underlies Arabidopsis leaf development. Plant Cell 20, 2293–2306 (2008).

7. K. Baerenfaller et al., Systems-based analysis of Arabidopsis leaf growth reveals adaptation to water deficit. Mol Syst Biol 8, 606 (2012).

8. A. Skirycz et al., Pause-and-stop: the effects of osmotic stress on cell proliferation during early leaf development in Arabidopsis and a role for ethylene signaling in cell cycle arrest. Plant Cell 23, 1876–1888 (2011).

9. J. Cao et al., The single-cell transcriptional landscape of mammalian organogenesis. Nature 566, 496–502 (2019).

10. C. Riou-Khamlichi, R. Huntley, A. Jacqmard, J. A. Murray, Cytokinin activation of Arabidopsis cell division through a D-type cyclin. Science 283, 1541–1544 (1999).

11. H. J. Kim, Y. H. Chiang, J. J. Kieber, G. E. Schaller, SCF(KMD) controls cytokinin signaling by regulating the degradation of type-B response regulators. Proc Natl Acad Sci U S A 110, 10028–10033 (2013).

12. Y. Jiang, G. Liang, S. Yang, D. Yu, Arabidopsis WRKY57 functions as a node of convergence for jasmonic acid- and auxin-mediated signaling in jasmonic acid-induced leaf senescence. Plant Cell 26, 230–245 (2014).

13. N. Gonzalez et al., Increased leaf size: different means to an end. Plant Physiol 153, 1261–1279 (2010).

14. Y. Li, L. Zheng, F. Corke, C. Smith, M. W. Bevan, Control of final seed and organ size by the DA1 gene family in Arabidopsis thaliana. Genes Dev 22, 1331–1336 (2008).

15. K. Morris et al., Salicylic acid has a role in regulating gene expression during leaf senescence. Plant J 23, 677–685 (2000).

16. D. N. Rate, J. V. Cuenca, G. R. Bowman, D. S. Guttman, J. T. Greenberg, The gain-of-function Arabidopsis acd6 mutant reveals novel regulation and function of the salicylic acid signaling pathway in controlling cell death, defenses, and cell growth. Plant Cell 11, 1695–1708 (1999).

17. B. J. DeYoung et al., The CLAVATA1-related BAM1, BAM2 and BAM3 receptor kinase-like proteins are required for meristem function in Arabidopsis. Plant J 45, 1-16 (2006).

18. S. Gan, R. M. Amasino, Inhibition of leaf senescence by autoregulated production of cytokinin. Science 270, 1986–1988 (1995).

19. J. Todd, D. Post-Beittenmiller, J. G. Jaworski, KCS1 encodes a fatty acid elongase 3-ketoacyl-CoA synthase affecting wax biosynthesis in Arabidopsis thaliana. Plant J 17, 119–130 (1999).

20. K. Prado et al., Regulation of Arabidopsis leaf hydraulics involves light-dependent phosphorylation of aquaporins in veins. Plant Cell 25, 1029–1039 (2013).

21. S. Gonzalez et al., Arabidopsis transcriptome responses to low water potential using high-throughput plate assays. Elife 12, (2024).

22. Z. Zhang, W. Li, X. Gao, M. Xu, Y. Guo, DEAR4, a Member of DREB/CBF Family, Positively Regulates Leaf Senescence and Response to Multiple Stressors in Arabidopsis thaliana. Front Plant Sci 11, 367 (2020).

23. S. J. Park et al., Ethylene responsive factor34 mediates stress-induced leaf senescence by regulating salt stress-responsive genes. Plant Cell Environ 45, 1719–1733 (2022).

24. A. Jain, G. T. Wilson, E. L. Connolly, The diverse roles of FRO family metalloreductases in iron and copper homeostasis. Front Plant Sci 5, 100 (2014).

25. I. Mukherjee, N. H. Campbell, J. S. Ash, E. L. Connolly, Expression profiling of the Arabidopsis ferric chelate reductase (FRO) gene family reveals differential regulation by iron and copper. Planta 223, 1178–1190 (2006).

26. G. Stefano et al., ER network homeostasis is critical for plant endosome streaming and endocytosis. Cell Discov 1, 15033 (2015).

27. A. Skylar, F. Sung, F. Hong, J. Chory, X. Wu, Metabolic sugar signal promotes Arabidopsis meristematic proliferation via G2. Dev Biol 351, 82–89 (2011).

28. L. M. Weaver, S. Gan, B. Quirino, R. M. Amasino, A comparison of the expression patterns of several senescence-associated genes in response to stress and hormone treatment. Plant Mol Biol 37, 455–469 (1998).

29. K. C. Dawei Dai, Jingwen Tao, Ben P. Williams, Aging drives a program of DNA methylation decay in plant organs. bioRxiv, (2024).

30. J. Swift, J. M. Alvarez, V. Araus, R. A. Gutierrez, G. M. Coruzzi, Nutrient dose-responsive transcriptome changes driven by Michaelis-Menten kinetics underlie plant growth rates. Proc Natl Acad Sci U S A 117, 12531–12540 (2020).

31. H. Claeys, S. Van Landeghem, M. Dubois, K. Maleux, D. Inze, What Is Stress? Dose-Response Effects in Commonly Used in Vitro Stress Assays. Plant Physiol 165, 519–527 (2014).

32. J. Swift, M. Adame, D. Tranchina, A. Henry, G. M. Coruzzi, Water impacts nutrient dose responses genome-wide to affect crop production. Nat Commun 10, 1374 (2019).

33. D. Martignago, A. Rico-Medina, D. Blasco-Escamez, J. B. Fontanet-Manzaneque, A. I. Cano-Delgado, Drought Resistance by Engineering Plant Tissue-Specific Responses. Front Plant Sci 10, 1676 (2019).

34. D. W. Lawlor, Genetic engineering to improve plant performance under drought: physiological evaluation of achievements, limitations, and possibilities. J Exp Bot 64, 83–108 (2013).

35. N. Illouz-Eliaz et al., Mutations in the tomato gibberellin receptors suppress xylem proliferation and reduce water loss under water-deficit conditions. J Exp Bot 71, 3603–3612 (2020).

36. C. Y. Cheng et al., Araport11: a complete reannotation of the Arabidopsis thaliana reference genome. Plant J 89, 789–804 (2017).

37. R. Satija, J. A. Farrell, D. Gennert, A. F. Schier, A. Regev, Spatial reconstruction of single-cell gene expression data. Nature Biotechnology 33, 495–U206 (2015).

38. J. Y. Kim et al., Distinct identities of leaf phloem cells revealed by single cell transcriptomics. Plant Cell 33, 511–530 (2021).

39. M. I. Love, W. Huber, S. Anders, Moderated estimation of fold change and dispersion for RNA-seq data with DESeq2. Genome Biology 15, (2014).

40. A. Sessions, D. Weigel, M. F. Yanofsky, The Arabidopsis thaliana MERISTEM LAYER 1 promoter specifies epidermal expression in meristems and young primordia. Plant J 20, 259–263 (1999).

